# Benchmarking the translational potential of spatial gene expression prediction from histology

**DOI:** 10.1101/2023.12.12.571251

**Authors:** Adam S. Chan, Chuhan Wang, Xiaohang Fu, Shila Ghazanfar, Jinman Kim, Ellis Patrick, Jean YH Yang

## Abstract

Spatial transcriptomics has enabled the quantification of gene expression at spatial coordinates, offering crucial insights into molecular underpinnings of diseases. In light of this, several methods predicting spatial gene expression from paired histology images have offered the opportunity of enhancing the utility of readily obtainable and cost-effective haematoxylin-and-eosin-stained histology images. To this end, we conducted a comprehensive benchmarking study encompassing six developed methods. These methods were reproduced and evaluated using HER2-positive breast tumour and human cutaneous squamous cell carcinoma datasets, followed by external validation using The Cancer Genome Atlas data. Our evaluation incorporates diverse metrics which capture the performance of predicted gene expression, model generalisability, translational potential, usability and computational efficiency of each method. Our findings demonstrate the capacity of methods to spatial gene expression from histology and highlight key areas that can be addressed to support the advancement of this emerging field.

## Introduction

Spatially resolved transcriptomics data (ST) capture the spatial organisation and heterogeneity of genes in tissues at high resolution, fundamentally transforming the way we study and understand biological processes. It has enabled the identification of key genes and pathways involved in intricate mechanisms underlying diverse biological processes and complex cellular phenomena ^1–4^. While this new ST technology holds promise for producing new insights, it remains costly, preventing its direct adoption into routine clinical use at this stage. In contrast, haematoxylin-and-eosin-stained (H&E) histopathology images are more cost-effective and are routinely used in clinics. Thus, if we can leverage modern machine learning together with information from ST technologies to predict in-silico spatial profiles of gene expression values from paired histology images of the same tissue, we could enhance information gain from existing H&E images. This would facilitate large-scale examinations of spatial gene expression variations and lead to the discovery of novel biomarkers and potential therapeutic targets for complex diseases.

To date, several approaches have been developed to predict in-silico spatially resolved gene expression (SGE) patterns using H&E data alone (**Figure 1a**) (see Supplementary Table 1) ^5–18^. Among these methods, Convolutional Neural Networks (CNN) and Transformers are commonly selected architectures for extracting local and global 2D vision features around each sequenced spot from corresponding histology image patches. Some methods further implemented extra components, including super-resolution enhancement of SGE ^9,18,19^ and Graph Neural Networks (GNN) ^8,12,16,18^ to capture neighbourhood relationships between adjacent spots. These learned features are then used to predict SGE that enable image-based screening for molecular biomarkers with spatial variation. As these methods were recently published in the last three years and continuously evolving, their performance has not yet been comprehensively benchmarked. While there exist general reviews on various aspects of spatial transcriptomics analysis ^20–22^, they have only surveyed and categorised methods predicting SGE from H&E without any benchmarking.

**Figure 1:**
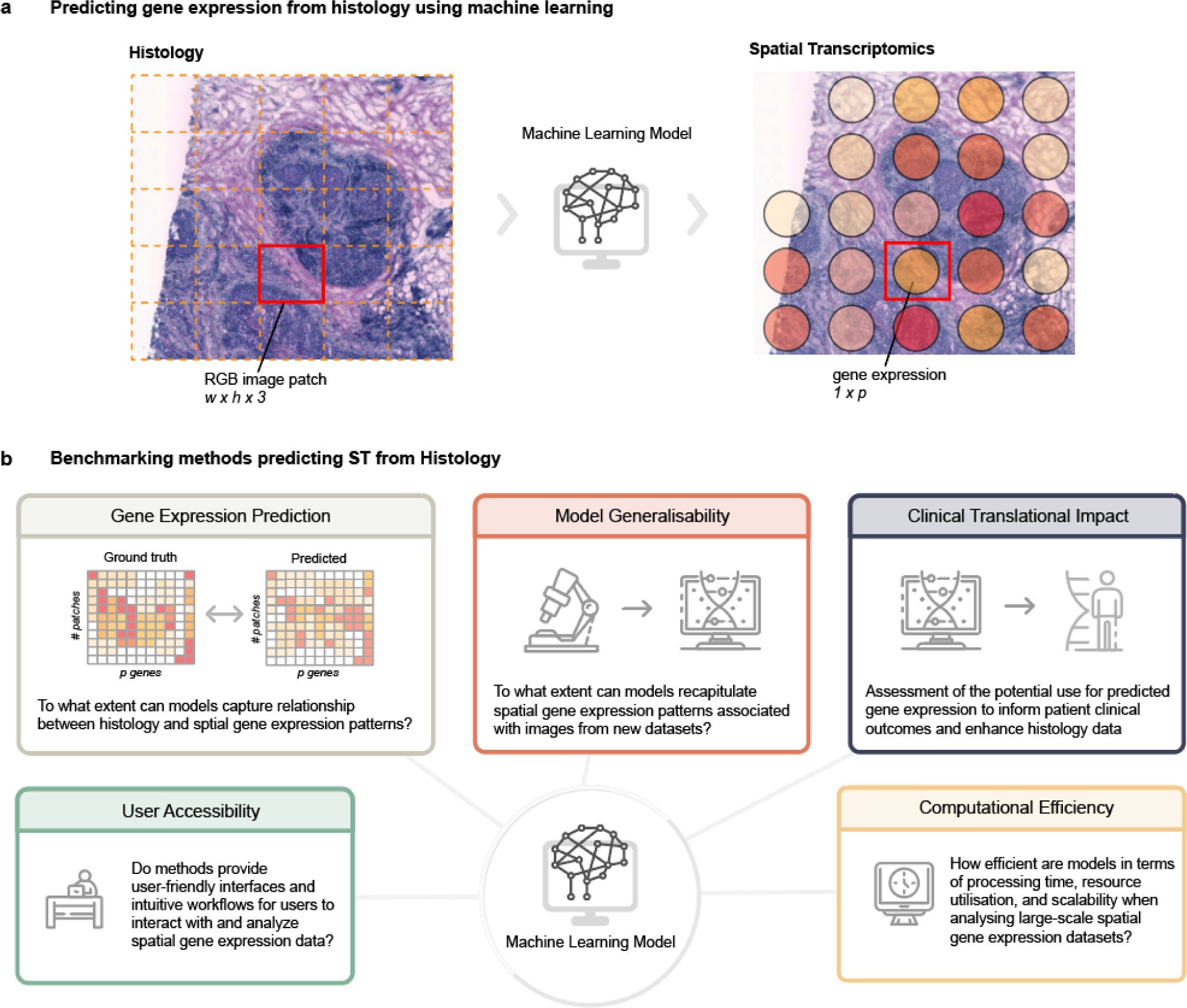
Overview of of the benchmarking process and the give key aspects of evaluation (a) Illustration of the machine learning task of predicting spatial gene expression from H&E images. (b) Overview of the evaluation categories used in the benchmarking of each method.

Evaluation studies that are presented when proposing a new method are limited in nature and often not independent. Among the current literature, methods for predicting SGE from H&E have been mainly assessed and evaluated for their efficacy by their developers. Evidently, different evaluation frameworks have been used by different developers to assess various aspects of their methods. For instance, ST-Net ^5^, DeepSpaCE ^9^ and DeepPT ^23^ evaluated the performance of their models on external The Cancer Genome Atlas (TCGA) data to determine the generalisability of their models. Hist2ST ^8^ performed an ablation study to assess the contributions of different components of their deep learning architecture. BrST-Net ^17^ directly compared 10 state-of-the-art CNN and transformer architectures to determine the effectiveness of the models. Overall, there is no consistent and detailed evaluation framework comparing the performance of these approaches in predicting in-silico SGE, mainly due to the fact that this field is still in its early development, data availability and the complexity of method implementations. Consequently, we lack a comprehensive understanding of the performance, stability and the broader applicability of individual methods. In addition to this there is a lack of critical evaluation regarding how different characteristics of H&E images can affect the accuracy of spatial SGE predictions, especially on its impact on clinical translation.

To this end, we provide the first benchmark and review of methods predicting SGE from H&E histology images. We compared the performance of six methods over 28 metrics broadly grouped into five categories. Several metrics on top of the standard performance measure of Pearson correlation are employed for understanding how well methods are able to recapitulate SGE in spatial transcriptomics data. Metrics also include performance measures stratified by gene characteristics to evaluate each method under different scenarios. With this emerging frontier of predicting SGE from H&E providing exciting potential for clinical usage, our benchmark study provides a vital assessment of the limitations and generalisability of their translational potential. Ultimately, our study aims to provide guidance for new users of these methods and highlight current challenges for developers aiming to advance methodologies.

## Results

### Benchmarking *in silico* spatial gene expression prediction from H&E histology image

To assess current methods of predicting gene expression from histology images, we have developed a benchmarking framework across five key categories (**Figure 1b, Methods**). We employed a hierarchy of evaluation categories: (1) within image SGE prediction performance, (2) across studies model generalisability, evaluated by applying models trained on ST data to The Cancer Genome Atlas (TCGA) images to identify whether models were useful for enhancing existing H&E histology images; (3) clinical translational impact by predicting survival outcomes and canonical pathological regions via predicted SGE. In addition, we considered (4) usability of the methods encompassing code, documentation and the manuscript; (5) and the computational efficiency (**Figure 1b**). These categories provide the foundation for comprehending the accuracy and applicability of histology-based gene expression prediction methods.

We employed an evaluation strategy that utilised 28 metrics across all evaluation categories to reveal diverse characteristics of six methods predicting SGE from histology images (**Table 1**). Our findings highlighted that no single method emerged as the definitive top performer across all categories of evaluation. HisToGene, DeepSpaCE and Hist2ST demonstrated notable performance in model generalisability and usability (**Figure 2a**), while DeepPT exhibited the best gene expression prediction accuracy, despite limited code availability at the time of assessment. Overall, the usability, model generalisability and clinical translation impact categories presented the most room for improvement across all methods, suggesting potential areas for further development within this field.

**Figure 2:**
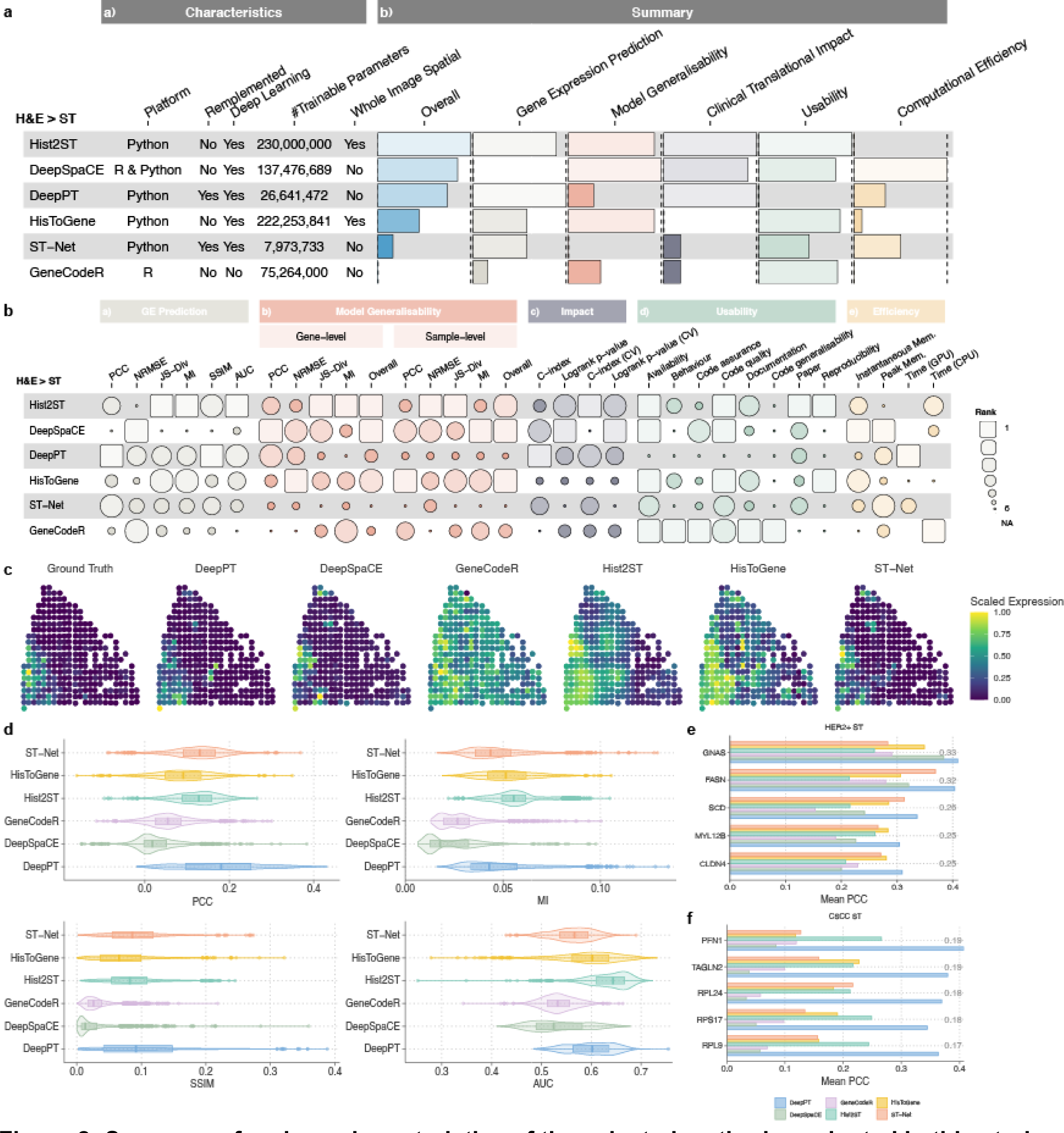
Summary of various characteristics of the selected methods evaluated in this study and their overall evaluation results. (a) Summary heatmap of methods predicting spatial gene expression from H&E images highlighting key characteristics, and ranking their performances under each evaluation category. (b) Detailed heatmap of rankings of each method under each evaluation metric grouped by category. (c) Spatial plots of FASN expression for one sample in the HER2+ dataset. (d) Boxplot/violin plots of the average PCC, MI, SSIM and AUC between the ground truth gene expression and predicted gene expression. Metrics measured from the test fold of a 4-fold CV, averaged over each gene across HER2+ and CSCC ST datasets. (e) Top five correlated genes in the HER2+ ST and (f) CSCC dataset. Grey number represents the correlation of the test set predictions averaged over each method.

**Table 1:** Architecture characteristics of methods benchmarked.

| Model | Platform | Deep Learning Model | Architecture Characteristics |  |  |  |
| --- | --- | --- | --- | --- | --- | --- |
|  |  |  | Local Features (one spot)<sup>a</sup> | Local + Global Features (spot-neighborhood relations)<sup>b</sup> | Global Features (spot-spatial relations)<sup>c</sup> | Trainable Parameters (in HER2+ ST) |
| ST-Net <sup>5</sup> | Python | Yes | Pretrained DenseNet 121 | NA | NA | 8M |
| HisToGene <sup>6</sup> | Python | Yes | ViT | Super Resolution | NA | 222M |
| DeepPT <sup>7</sup> | Python | Yes | Pretrained ResNet50 + Autoencoder + MLP | NA | NA | 26M |
| Hist2ST <sup>8</sup> | Python | Yes | Convmixer | GNN - GraphSAGE | Transformer | 230M |
| DeepSpaCE <sub>60</sub> | R & Python | Yes | VGG16 | Super Resolution | NA | 137M |
| GeneCodeR <sub>10</sub> | R | No | NA | NA | NA | 75M |

<sup>a</sup> local features within a patch
<sup>b</sup> neighbouring global/contextual features within a section of histology image
<sup>c</sup> global/contextual features within the whole histology image

### All methods are able to capture biologically relevant gene patterns from tissue images

The use of several evaluation metrics identified methods that are better performing for predicting SGE from histology. After models were trained to predict SGE from histology, we compared the predicted SGE to the ground truth SGE in hold-out test images from cross-validation (**Figure 2c**). Across the HER2+ and cutaneous squamous cell carcinoma (CSCC) ST datasets, we measured the relative performance based on Pearson Correlation Coefficient (PCC), Mutual Information (MI), Structural Similarity Index (SSIM) and Area Under the Curve (AUC) (**Table 2**) for each method (**Figure 2d**).

**Table 2:**
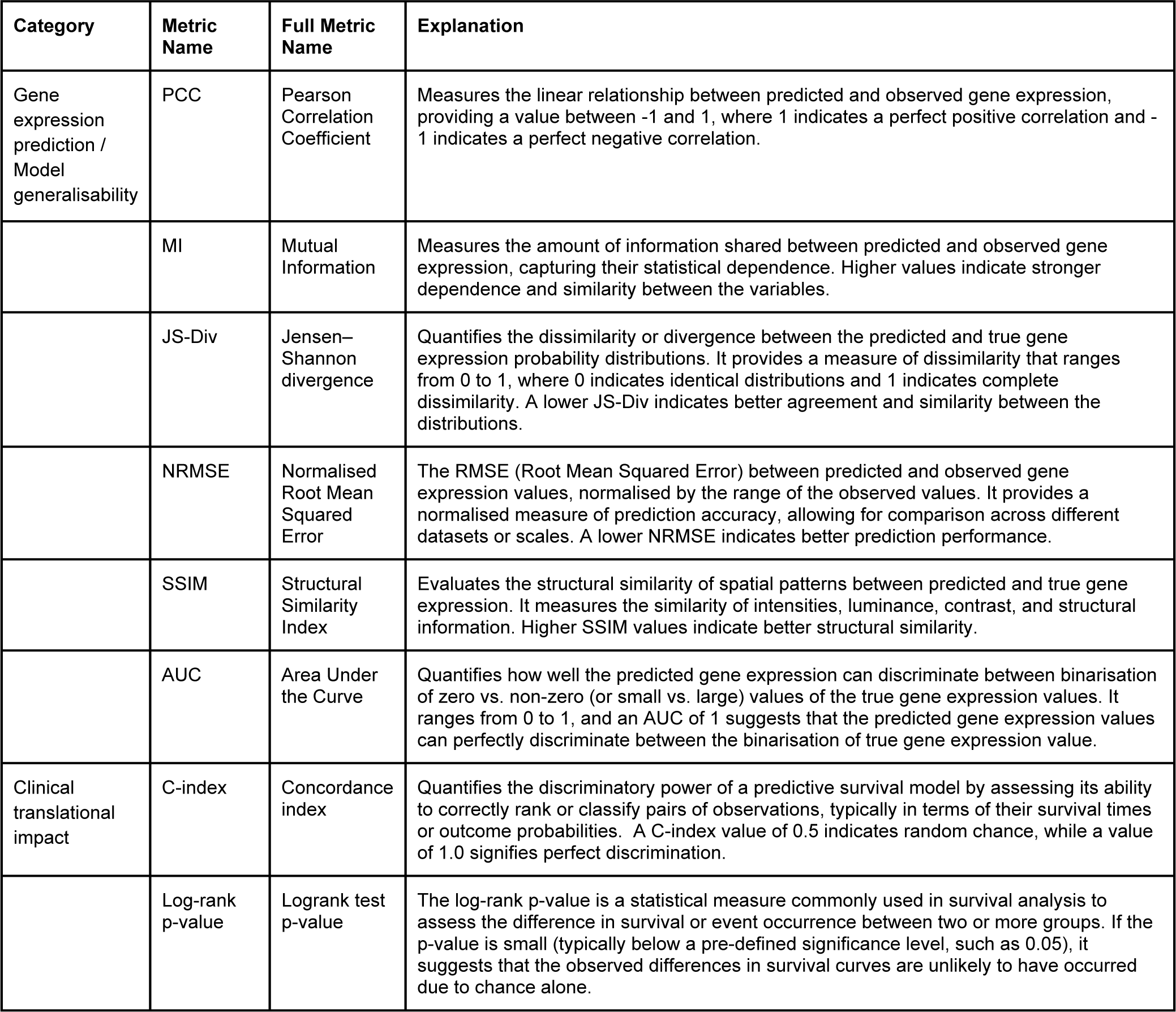
Evaluation metric used and their interpretations.

DeepPT had the highest average correlation of (PCC = 0.11; SSIM of 0.07) across all genes, suggesting that its predicted GE and spatial patterns were more aligned to the ground truth GE patterns than other methods. On the other hand, Hist2ST had the highest average MI of 0.61 and AUC of 0.61, suggesting the predicted GE displayed some trend with the ground truth and was able to distinguish between zero and non-zero expressions better than other methods. DeepPT, characterised by a simpler CNN architecture (**Table 1**), demonstrated distinct predictive capabilities compared to Hist2ST, which additionally captures the global spatial features on the whole image slide using the learned features from each spot. The aggregation of the SGE prediction metrics ranked DeepPT as the best performer for SGE prediction, followed by Hist2ST, HistToGene, ST-Net, GeneCodeR and DeepSpaCE.

Subsequently we examined if these methods captured genes specific to the tissue types in the data. Here, we assessed the capacity for methods to capture tissue relevant GE by examining the biological relevance of the top correlated genes. Here, DeepPT observed the highest correlations in genes GNAS (r = 0.41) and FASN (r = 0.4) in HER2+ ST dataset (**Figure 2e**) and PFN1 (r = 0.41) in CSCC ST dataset (**Figure 2f**). In the context of HER2+ breast cancer, FASN is known as a hallmark feature observed early in most human carcinomas and precursor lesions which has been associated with adverse patient outcomes and therapeutic resistance ^24^. This gene is one of the highest predicted genes (r = 0.32 across HER2+ and CSCC) identified across all methods (**Figure 2e**) and is aligned with key cancer-related molecular pathways ^24^. Additionally, MYL12B, with an average correlation of 0.26, is known to be involved in the regulation of cell morphology with links to cancer progression ^25^. In the CSCC dataset, LMNA (r = 0.17), has been shown to have increased expression in skin cancer ^26^. These findings indicate that all methods, despite the relatively low average correlation, were able to capture biologically relevant gene patterns from tissue images (see Discussion).

To further investigate performance, we sought to ascertain how well each approach extracted biologically relevant insights from spatial transcriptomics data, whilst maintaining a focus on excluding the influence of data noise. We assessed this by calculating PCC and SSIM values within highly variable genes (HVG) only (**Figure 3a**) and compared to the performance across all genes. Under HVGs only, most methods increase in correlation (DeepPT, GeneCodeR, p < 0.05) or SSIM (DeepPT, DeepSpaCE, GeneCodeR, Hist2ST, ST-Net, p < 0.05) compared to using all genes and we believe this provides a more meaningful evaluation of methods. Furthermore, we delved into the capacity of the methods to discriminate between different levels of GE, employing several binary thresholds (**Figure 3b**). Our findings revealed that the predicted SGE across all methods was able to distinguish between ground truth GE counts greater than 10 and less than 10 with a mean AUC=0.66. This outperformed the ability to predict between non-zero and zero GE where mean AUC=0.58, suggesting that methods were more capable of distinguishing between higher and lower values of GE. We also found that all methods had a negative correlation between the predicted vs ground truth correlation and the percentage of zeros for each gene (p < 2 × 10^−5^), which was more notable for DeepPT, GeneCodeR and ST-Net (**Supplementary** Figure 3). For the strongest performing method in GE prediction, DeepPT, the performance was noticeably affected by the variance, absolute expression and percentage of zeros of the genes it was trained on. Thus we observed that some methods perform better when noisy genes are removed from the analyses.

**Figure 3:**
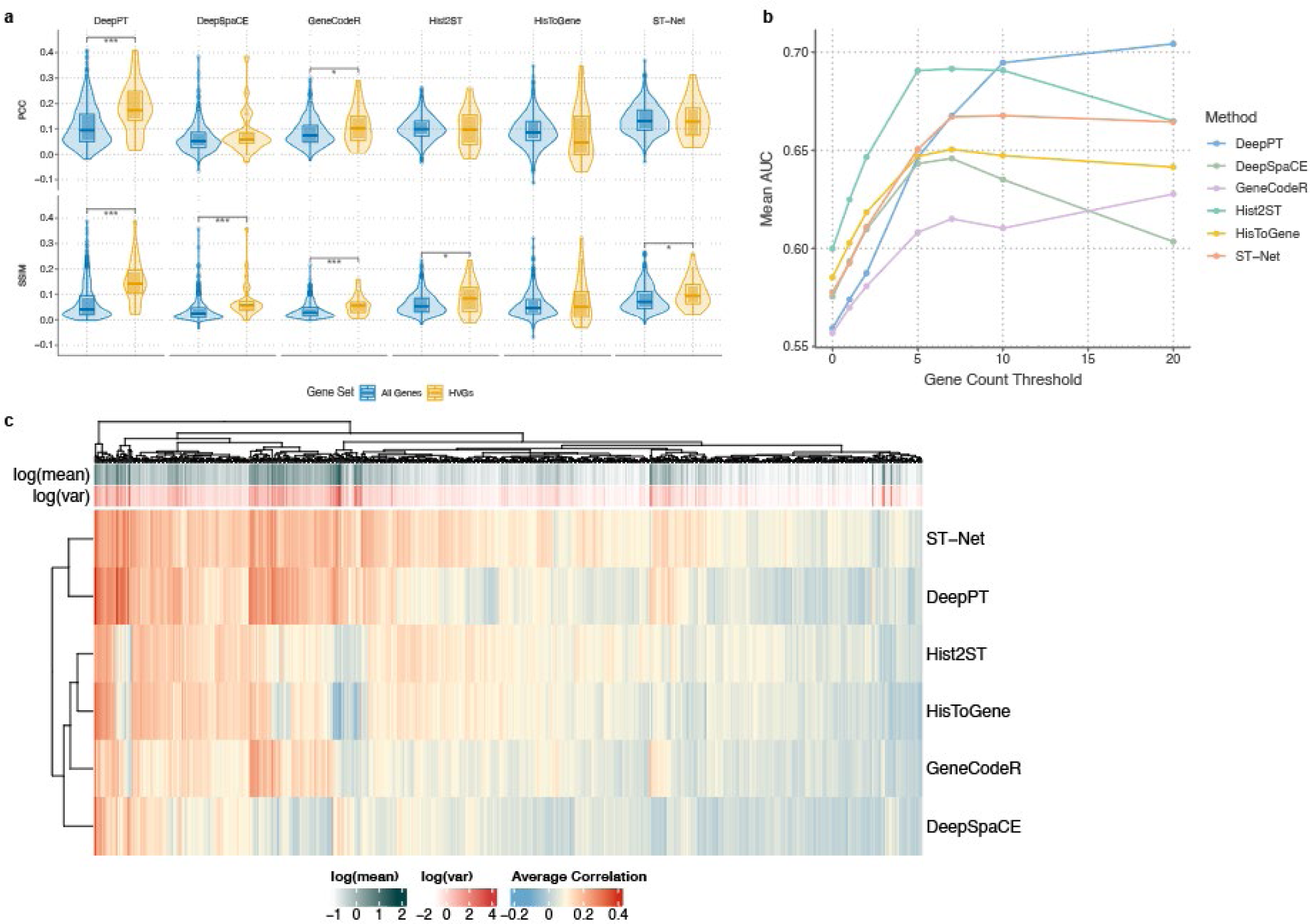
Investigation of correlations and performance variation among methods in HER2+ dataset (a) PCC and SSIM violin and boxplots for each method in HER2+ dataset for all genes as well as for HVGs only. Significance between HVGs and all genes are calculated using Wilcoxon rank-sum test (The significance levels were defined as °p<0.1, *p < 0.05, **p < 0.01, ***p < 0.001). (b) Plot of the AUC (averaged over each gene) of the predicted gene expression distinguishing a binarisation of the ground truth gene expression. Ground truth is binarised according to whether the value was greater than or equal to several thresholds (x-axis). (c) Heatmap of average correlation of each gene in HER2+ ST dataset and each method. The log of the mean and variance of each gene are coloured above the heatmap.

### Characteristics of methods play a large role in gene-specific performance

We set out to uncover the role of varying deep learning architectures in gene-specific prediction performance and the extent to which these structures shaped our results. In **Figure 3c**, we plotted the average correlation of each gene over each method in the HER2+ ST dataset. We then employed hierarchical clustering on the correlations from each gene and method to identify the similarities between method performance based on characteristics of their deep learning architectures. Notably, we observed that DeepPT and ST-Net clustered together, a finding consistent with their shared utilisation of pre-trained CNNs focusing on single image patches only (**Table 1**). Hist2ST and HisToGene formed another distinct cluster, attributed to their emphasis on global interactions and their models taking whole slide images as input as opposed to just image patches in the other methods.

Hence, the similarities in method performance aligned with the inherent structural characteristics of these methods.

Furthermore, we aimed to determine differences in gene-specific performance due to different data processing between methods. A substantial proportion of genes displayed positive correlations across all tested methods (**Figure 3c**), highlighting common genes that were more generally well predicted such as MYL12B, SCD and GNAS. We also identified a subset of genes that exhibited strong performance in DeepPT and ST-Net but displayed diminished performance in other models such as C3 and BGN. This occurred due to an observed reduction in correlation between the original GE data and the normalised data used for model training, from normalisation techniques specific to the Hist2ST and HisToGene methods (**Supplementary** Figure 4). These correlation findings highlight an interplay between methods input processing and SGE gene-specific prediction performance.

### Comparing the translational potential across methods using TCGA-BRCA data

We investigated the translational potential of predicted SGE for diagnostics in pathology as one of the downstream applications of predicting SGE from H&E. We evaluated this from three perspectives, (1) robustness to input quality; (2) generalisability to an independent cohort; and (3) utility in survival prognosis. Firstly, we assessed whether H&E image quality impacted the accuracy of prediction. For the TCGA-BRCA stage I breast cancer (BC) H&E images, we first calculated a wide range of H&E quality control (QC) metrics (**Supplementary** Figure 5) using HistoQC ^27^. Next, we applied the methods pretrained on the HER2+ ST data to predict bulk RNA-Seq GE patterns for these images.

We used the patient-level correlation as a measure of prediction performance by calculating the correlation between predicted GE pseudobulk and bulk GE for each image. We then measured the correlation between the first principal component of the H&E QC metrics and prediction performance over each image and method. Previous literature ^28^ had shown that the quality of H&E impacts segmentation performance and so we expect that a robust method will exhibit minimal dependence between image quality and the SGE prediction performance. DeepPT showed an undesirably high correlation (r = 0.53) between image quality and prediction performance (**Figure 4a**). Meanwhile, HisToGene (r = 0.05) and Hist2ST (r = 0.19) had little to no dependence on the obtained QC metrics, indicating their robustness to image quality. HisToGene and Hist2ST used normalisation in their workflow, which indicates that input processing may help reduce dependence on image quality.

**Figure 4:**
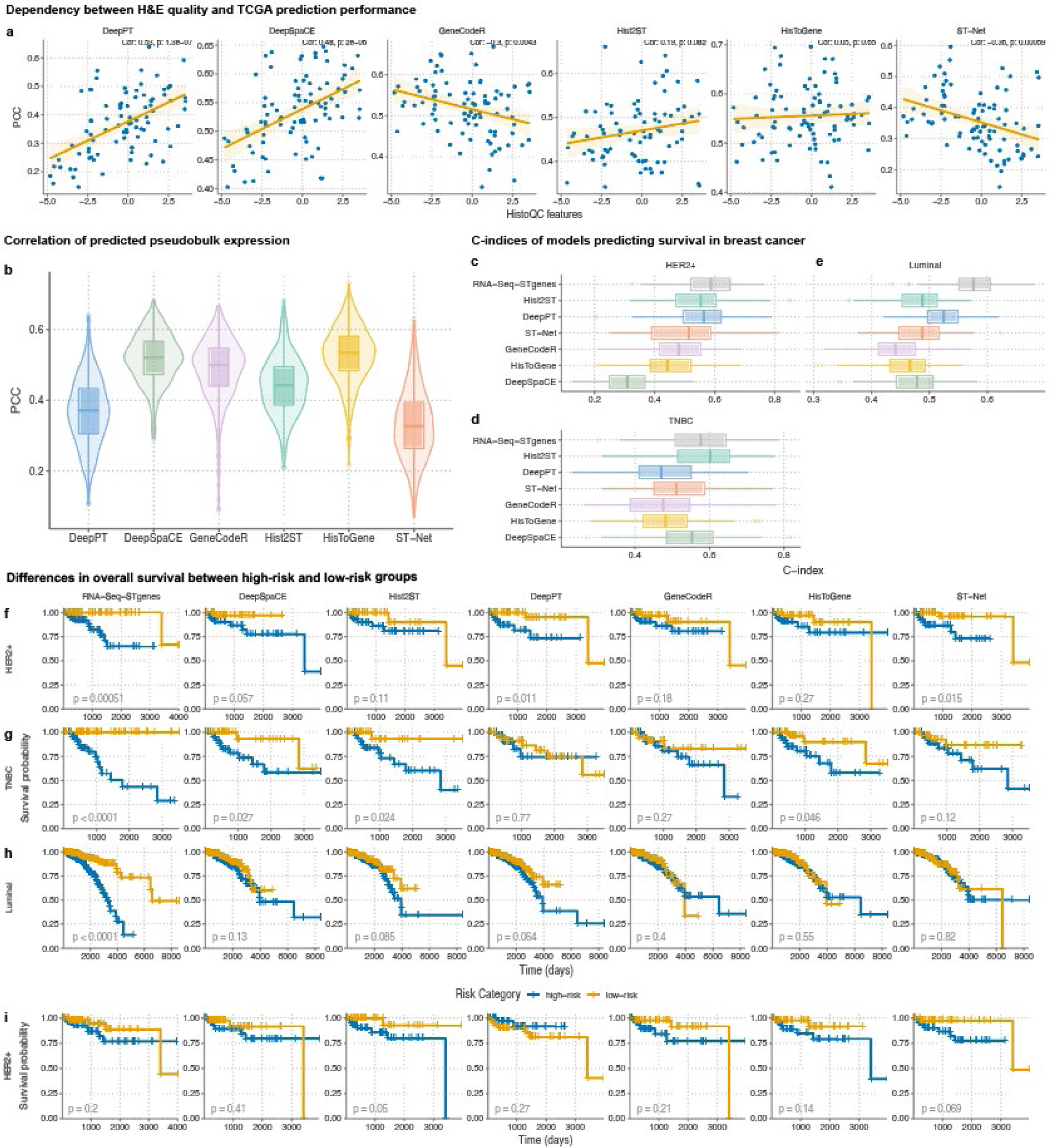
Evaluation on translational potential of gene expression predictions (a) Scatterplot of patient-level correlations between predicted pseudobulk gene expression and RNA- Seq bulk gene expression vs. first principal component of calculated H&E quality control metrics. H&E QC metrics were calculated on stage I breast cancer patients. Line of best fit plotted along with Pearson correlation and associated p-value from a correlation test. (b) Boxplot of patient-level correlations between predicted pseudobulk gene expression and RNA-Seq bulk gene expression over all analysed TCGA-BRCA images split by method. Models were trained on the HER2+ ST dataset, selecting the model from the 4-fold CV with the best test set performance. (c-e) C-indices of multivariate cox regression models predicting survival of TCGA-BRCA patients, using RNA-Seq bulk, RNA-Seq bulk using only genes present in HER2+ ST dataset, and the predicted pseudobulk from each method. C-indices were calculated from the test sets of a 3-fold CV with 100 repeats trained within (c) HER2+, (d) luminal and (e) TNBC breast cancer clinical subtypes. (f-i) Kaplan-Meier curves for patients split into high and low risk groups by the median risk prediction of the multivariate cox regression models for each method and (f) HER2+, (g) luminal and (h) TNBC breast cancer subtypes. Models were trained on all patients and then the predictions of the training patients were used. The p- value represents the result of the logrank test for assessing the statistical significance of differences in survival between the groups. (i) Kaplan-Meier curves constructed using the average risk prediction from a 3-fold CV with 100 repeats within the HER2+ breast cancer subtype.

Hence, it is crucial to take image quality into account during method development, as it can have unintended effects on performance.

We further assessed the generalisability of all methods by determining whether methods trained on ST data could feasibly be used on an independent set of H&E images. Using the predicted pseudobulk GE from each method and the bulk GE associated with the TCGA-BRCA H&E images, we calculated the patient-level correlations as defined above. The patient-level correlations averaged across methods were then used to compare the translational performance (**Figure 4b**). The methods HisToGene (mean = 0.53) and DeepSpaCE (mean = 0.54) performed the best in this section. These methods were followed by GeneCodeR (mean = 0.49), Hist2ST (mean = 0.44), DeepPT (mean = 0.37) and ST-Net (mean = 0.33), suggesting that all methods were able to capture the overall trends and variations in GE for the majority of patients. The positive patient-level correlations suggest that the methods have utility in providing GE information for histology images that do not necessarily have associated spatial GE data.

We then examined the downstream utility of predicted pseudobulk GE from H&E in breast cancer survival analysis in the TCGA-BRCA images. For each method, we built multivariate Cox regression models from the pseudobulk GE to predict survival for three subtypes of breast cancer (BC): HER2+ BC, triple-negative breast cancer (TNBC) and luminal BC (as previously shown in ^29^). We compared these models against models built using the bulk RNA-seq (baseline) data using average C-index from a 3-fold cross validation (CV) with 100 repeats. In HER2+ BC patients, DeepPT had the highest average C-index=0.55 compared to 0.58 for the RNA-Seq (**Figure 4c**). For TNBC, Hist2ST had the highest average C-index of 0.58, compared to 0.57 for RNA-Seq (**Figure 4d**). For luminal BC, DeepPT had the highest C-index of 0.52 compared to 0.58 for RNA-Seq (**Figure 4e**). Across all BC subsets, Hist2ST had the highest average C-index from evaluating using CV and DeepPT had the highest C-index from evaluating using the training data (**Supplementary** Figure 6). Subsequently, we binarised the risk model predictions into high and low-risk groups through using median prediction to split patients then constructed Kaplan-Meier (KM) survival curves. Following the analysis of Xu et al. (^29^), we were able to capture the same distinct KM curves for the RNA-Seq models in each BC subset of patients (*p* ≤ 5.1 × 10^−4^) using the risk predictions of the patients the models were trained on (**Figure 4f-h**). However, we also showed that under cross-validation the distinction between high and low-risk groups were no longer significant except for Hist2ST (*p* = 0.05) and ST-Net (*p* = 6.9 × 10^−2^) in the HER2+ subset (**Figure 4i**). Under cross-validation, the log-rank p-values along with the c-indices indicated that the predicted GE from the models together with the RNA-seq bulk data may not be fully prepared for clinical translation. Despite the limited performance in unseen test data, the relative strength of each method under clinical translation potential were still able to be compared and these results highlighted an area for further refinement of these models.

### Usability of methods remains a major challenge

We evaluated the user accessibility for each method through a broad survey assessing several aspects of usability. We used a scoring schema (**Figure 5**) based on existing tool quality and programming guidelines in literature and online ^30^. These aspects included availability of software, code quality, code assurance, documentation, behaviour, paper, reproducibility, and generalisability. Overall scores for usability were obtained after summing weighted scores over each aspect (**Supplementary Table 4**), with a maximum attainable score of 32. Hist2ST achieved the highest usability score (18.8), followed by GeneCodeR (15.9), DeepSpaCE (15.5), HisToGene (16.5), ST-Net (10.2), and then DeepPT (2, lowest due to not having any code available at time of assessment). The mean usability score was 15.4 (sd = 2.8) excluding DeepPT. The majority of methods had not been tested for robustness in their paper and we were not able to be run without significant adjustments to the code. In contrast, Hist2ST excelled in these areas. Along with Hist2ST, HisToGene also provided a relatively user-friendly tutorial. Aside from DeepPT, ST-Net had the lowest overall score due to issues with code execution and undocumented functions, necessitating our own implementation for benchmarking ^6^. While the scoring schema used was able to inform which methods were the most user accessible methods, and provide a perspective on how these methods compare to exemplary tools and programs. It is important to note that overall, there was a clear gap between the maximum usability score and the highest scoring method.

**Figure 5:**
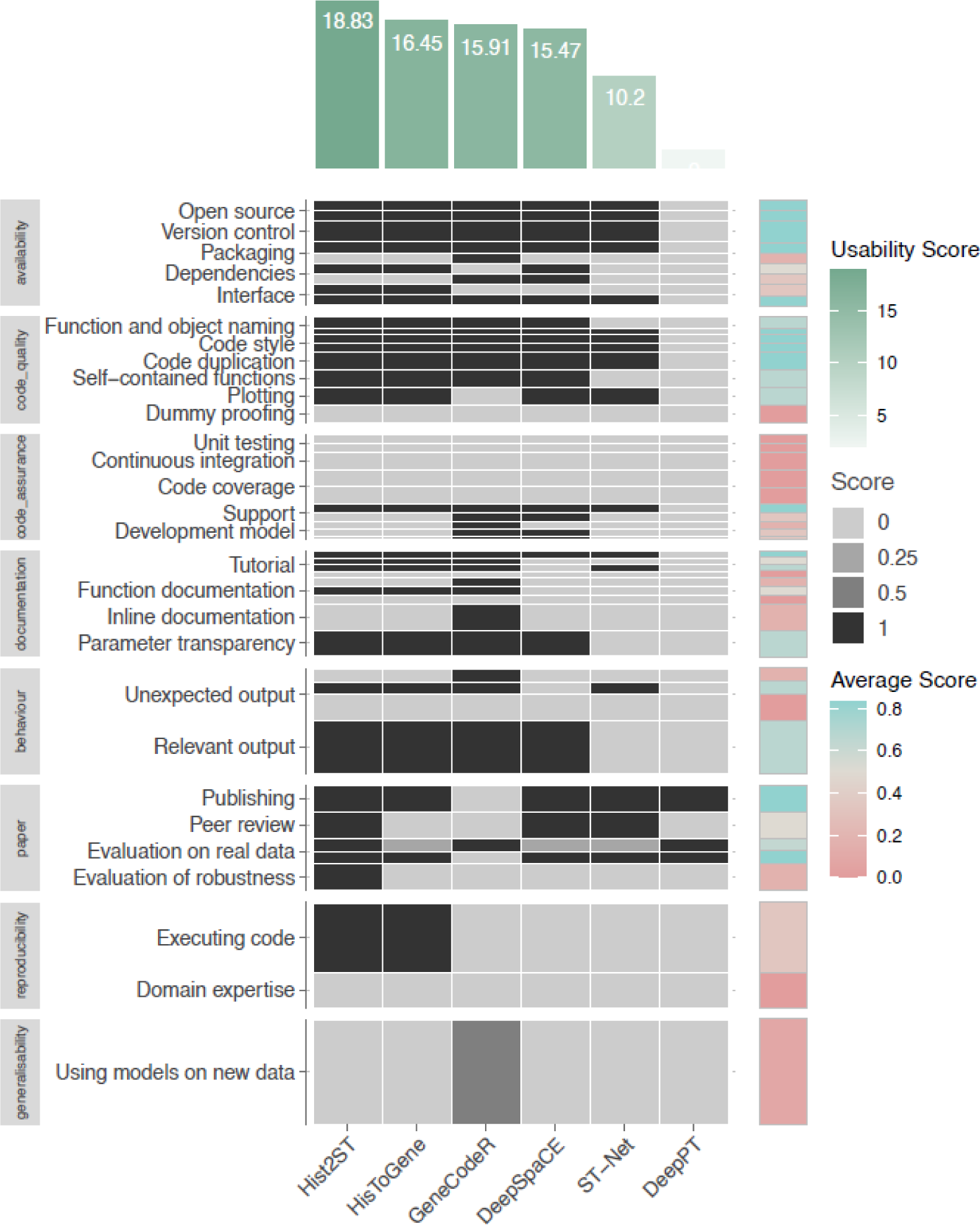
Usability score for the various methods. Heatmap showing the scores of each method under each usability scoring category. Height of each box represents the weight of each criteria contributing to the overall score plotted in the green bar plot on top. The blue-pink heatmap represents the row-wise average of the category scores over all methods.

We unpacked and examined the subcategories within the usability scores that are most needed for improvement. **Figure 5** shows a heatmap that highlighted subcategories of usability, with pink indicating weakness and blue indicating strength. The weakest usability subcategory was code assurance, which includes unit testing, support, development model, and continuous integration.

Whilst most methods were available on Github and had a support ticketing system, only GeneCodeR and DeepSpaCE created separate releases and branches for development and master code. All of the methods failed to incorporate any unit testing for the prevention of unexpected code behaviour with any updates. Key strengths included open-source code, coding consistency, exposed important parameters, and no unexpected files generated. Subcategories that require substantial improvement (average score = 0) included the need for knowledge of deep learning parameters and PyTorch syntax, unexpected warnings during compilation, undocumented function outputs, limited tutorials, absence of dummy-proofing, and often requiring users to write new code to apply models. Addressing these challenges would enhance overall usability, expanding the methods’ user base and research potential.

## Discussion

Here we presented a comprehensive benchmarking study of six developed methods for the prediction of spatial GE from H&E histology images. Our evaluation extends beyond standard performance measures such as correlation, by incorporating biologically informed metrics measuring HVGs and expression levels of genes. Furthermore, we tested the translational potential for these methods, examining model robustness to input quality, generalisability and the effectiveness of predicted genes by assessing their capabilities in survival prediction. Our study impartially evaluates the current state of methods that predict SGE from H&E histology images, with a particular focus on their potential to be applied in clinical contexts.

All evaluations to date strongly relied on correlation between observed and predicted gene expression as a quantitative measure to provide evidence of predictability of SGE from H&E. Our current benchmark prompts a discussion on whether correlation based on all genes should be the primary metric for evaluating methods. Most studies, including this benchmark, demonstrate that all methods consistently exhibited average correlations below 0.2, with a typical variability of around 0.2. It is important to note that we should not anticipate any correlation between two sets of white noise data, therefore genes with no expression, or expression at the limits of detection, should not have correlation between predicted and measured intensity. Hence, the proportion of low expressed or non-expression genes are high with the calculation of average masking the identification of biological meaningful concordance. As an alternative, we believe it is more meaningful to focus on differentially expressed or highly expressed genes, acknowledging that correlation is more meaningful among signal rather than noise and consequently, providing a more meaningful assessment of performance.

Following from the point above, since the average correlation between predicted expression and measured expression was low (0.2), predicted expression methods did not demonstrate readiness for use in clinical applications based on our TCGA study. Although we obtained higher patient-level correlations predicting bulk GE (mean = 0.45), there was a lack of demonstrative capacity in survival analysis via the predicted pseudobulk GE, as the methods were not able to stratify patients into high-risk and low-risk groups well and achieved poor C-indices lower than 0.6 via cross validation. This provides impetus to explore the suitability of deep learning architectures, the effect of normalisation techniques on predicted GE, and the impact of the multivariate cox regression model on translational ability to clinical applications. It would also be meaningful to understand how these methods perform in comparison to classification from H&E image features alone and quantify how much additional value spatial GE information can add to images for clinical prediction tasks. Notwithstanding, the comparison of survival prediction performance, which existing methods have not explicitly explored, can serve as a useful measure for guiding future method development.

Our overall observation is that the most complex deep learning architectures involving more components were not superior in either SGE prediction or in translational potential categories. This is surprising given the promise of newer deep learning techniques that have consistently demonstrated strong performance in other fields ^31–34^. Upon closer examination of different deep learning architectures, we found that models effectively capturing both local and global features within single-spot patches, as exemplified by DeepPT, outperform models attempting to incorporate additional spatial information from neighbouring spots or the entire image. This observation suggests that GE patterns in adjacent spots may not necessarily be strongly or consistently correlated. This may not be too surprising given that the spots in our evaluation data are around 150-200 μm apart from centre-to-centre. Thus, including global information from the entire image slide may introduce noise since there is a large number of genes and some genes are not expressed across all spots. Interestingly, transformer-based models demonstrated better risk group stratification, suggesting that their ability to capture various histology characteristics could be more closely related to downstream clinical outcomes than spatial GE patterns. Therefore, it is essential to delve deeper into the biological correspondence between H&E images, GE and corresponding clinical outcomes.

As methods often contain many components and functions, it is becoming vital to consider usability to minimise overhead and enhance the community’s capacity in using and testing methods. We provide here some suggestions for improving the usability of methods that predict ST from H&E images. As a minimum, the source code should be provided in an open source repository ^35–37^ such as GitHub, which was the case for five out of six of the methods benchmarked. Open source repositories should foster community support through issue tracking and participation in forums. Furthermore, there is typically high variability in the code organisation and software dependencies used. The documentation should state the software dependencies ^38,39^, especially the versions of Python/R and other packages to ensure that there are no conflicts or differences in behaviour. System resources, including CPU and non-standard hardware (e.g., GPU) requirements, should also be provided to ensure sufficient computational resources in the system ^37,40^. Instructions on installation and running of the scripts should be provided in the README file. Often, the repository contains multiple scripts that perform a series of processes with non intuitive ordering. The order in which the scripts should be run should be given, either stated in the README file, by numbering the scripts, or in a sample script that runs the other scripts ^37,41^. In addition, a demo dataset in the format required by the source code will help with ensuring data from various other ST data formats are correctly preprocessed ^39^. Finally, a brief description of outputs (either a sample output or list of files) will help with verifying that the code has run as expected.

Our benchmark elucidated the comparative results of the SGE prediction methods by employing a hierarchy of evaluation metrics. These results uncovered the importance of accounting for SGE attributes, histology image quality, as well as the choice of deep learning architecture in SGE prediction performance. Our findings demonstrate promising practical applications of these models in clinical contexts while also highlighting areas for further development, particularly in predicting patient survival using histology alone. Additionally we identified the main challenges for the widespread use of the methods related to their usability. We believe that this evaluation can provide a holistic, comprehensive framework for evaluating developed methods and provide guidance for future developments in this continually evolving field. Continued advancements in methods for predicting SGE from H&E histology can create opportunities for profound insights into complex spatial relationships and the potential to revolutionise disease risk diagnosis for patients.

## Materials and Methods

### Benchmarked methods selection criteria

We considered all methods published and in pre-print that predicted spatially resolved transcriptomics data using histology images up until June 2023 (**Supplementary Table 1**). As there were several methods which were recently developed, not all methods included publicly available code at the time of investigation (e.g. TCGN ^16^, BrST-Net ^17^, NSL ^15^, TransformerST ^18^ and STimage ^42^) were excluded as a result. BLEEP ^14^ was unable to be run due to lack of tutorials, and EGN ^11^ was run however only the training set produced reasonable correlations and thus was excluded from our results. Methods that performed joint modelling of H&E (e.g. Cottrazm ^43^, starfysh ^44^, and xfuse ^13^) were also excluded as we were only investigating the applicability of H&E to ST. In addition, methods predicting or incorporating scRNA-seq (e.g. SCHAF ^45^, St2cell ^46^ and ^47^) or bulk RNA-seq (e.g. HE2RNA ^48^ and ISG ^49^) were excluded. Although we excluded all methods predicting bulk gene expression from histology, we adapted the architecture from DeepPT ^23^ for predicting spatial transcriptomics. This was done as the model architecture was relatively clear to reproduce and the authors had also aimed to predict clinical outcomes using predicted SGE from their method, which was one of our evaluation categories of interest.

### Datasets and preprocessing

The same publicly available Spatial transcriptomics (ST) data that were employed in previous studies were used for benchmarking. In addition, H&E images matched with bulk gene expression profiles from TCGA were also used.

### Human HER2-positive breast tumour ^50^

The HER2+ dataset was measured using ST to investigate SGE in HER2-positive breast tumours, from which the original investigators discovered shared gene signatures for immune and tumour processes. The dataset consists of 36 samples of HER2-positive breast tissue sections from 8 patients.

For the histology images, 224 × 224 pixel patches were extracted around each sequencing spot. For the SGE data of each tissue section, the top 1,000 highly variable genes for each section were considered and those genes that were expressed in less than 1,000 spots across all tissue sections were excluded. This resulted in 785 genes for the training of all models.

### Human cutaneous squamous cell carcinoma ^51^

The CSCC dataset was measured using ST to understand cellular composition and architecture of cutaneous squamous cell carcinoma. The original investigators discovered shared gene signatures for immune and tumour processes. Whilst the original dataset contained single-cell RNA sequencing along with ST and multiplexed ion beam imaging, we analysed only the ST data. This ST data consists of 12 samples from 4 patients.

For the histology images, 224 × 224 pixel patches were extracted around each sequencing spot. For the gene expression data of each tissue section, the top 1,000 highly variable genes for each section were considered and those genes that were expressed in less than 1,000 spots across all tissue sections were excluded. This resulted in 977 genes for the training of all models.

### TCGA Breast Invasive Carcinoma (TCGA-BRCA)

Paired RNA-seq bulk gene expression and H&E histology images from The Cancer Genome Atlas Network ^52^ were used to evaluate model generalisability. This dataset was chosen as the predictions on these H&E images could be evaluated through the matched gene expression, and since the tissue type is the same as the data the models were trained on (HER2+).

TCGA-BRCA data was downloaded using the TCGAbiolinks package ^53^ version 2.29.6. RNA-Seq data was obtained by following query options: project = “TCGA-BRCA”, data.category = ”Transcriptome Profiling”, data.type = ”Gene Expression Quantification”, experimental.strategy = “RNA-Seq”, and workflow.type=“STAR - Counts”. Histology images were obtained by the following query: project = “TCGA-BRCA”, data.category = “Biospecimen”, data.type = “Slide Image”, and experimental.strategy = “Diagnostic Slide”. We only considered images that had one associated RNA-Seq entry and were either stage I, stage III, or stage IIA due to storage limitations. Overall we considered 671 TCGA images. Images were read using the openslide package, and converted to jpg from svs. Images were then split into 224 x 224 pixel patches and filtered out if the sum of the rgb values were greater (or more white) than a pure rgb(220,220,220) image. Further clinical information containing variables to define breast cancer subtypes for samples were downloaded from Broad GDAC Firehose (http://gdac.broadinstitute.org/).

For Hist2ST, the model fuses image patch embedding with spot coordinate embedding; a few TCGA images have more than 2048 spots exceeding the number of embedding dimensions. Therefore, the selected TCGA images for Hist2ST were cropped into 9 large patches before further splitted into 224x224 patches.

### Setting used and practical implementation of existing methods

#### GeneCodeR

GeneCodeR was the only method that did not use deep learning for its model. The generative encoder was set up as the following relation between the histology images (*H*) and SGE values (*E*):

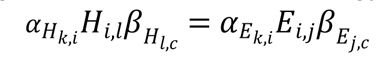

where the *α* and *β* are matrices that transform the sample and features across the datasets into the same space, *k* and *c* are the dimensions of the reduced dimensional encoding for the samples and features respectively, *i* is the number of samples, *j* is the number of genes and *l* is the number image dimensions.

The following hyper-parameters were used for the coordinate descent updating algorithm: the dimensions of the encoding space for the samples (*i_dim*) of *k*=500, the dimensions of the encoding space for the features (*j_dim*) of *c*=500, initialisation for the *alpha_sample* = “rnorm”, initialisation for the *beta_sample* = “irlba”, max iterations of 20, tolerance of 5, learning rate of 0.1, and batch size of 500.

#### HisToGene

HisToGene, is a deep learning model for SGE prediction from histology images. HisToGene employs a Vision Transformer, a state-of-the-art method for image recognition to predict SGE and can also be used to enhance resolution of SGE. The SGE values used for training were the normalised UMI counts from each spot. The normalised counts were derived by taking the UMI count for each gene divided by the total UMI counts across all genes in that spot, multiplied by 1,000,000, and then transformed to a natural log scale.

The HisToGene model was then implemented using PyTorch with the following hyper-parameters: a learning rate of 10^−5^, the number of training epochs of 100, a drop-out ratio of 0.1, 8 Multi-Head Attention layers and 16 attention heads. The final model used was the model after all epochs, as per the HisToGene pipeline.

#### DeepSpaCE

The Deep learning model for Spatial gene Clusters and Expression, DeepSpaCE, was developed for the imputation of tissue sections and for the enhancement of resolution of SGE. A VGG16 deep CNN model architecture is used for SGE prediction and semi-supervised training is available in DeepSpaCE. The DeepSpaCE pipeline was originally developed to read data output from the Space Ranger pipeline (10x Genomics), so the code had to be configured to read the spatial transcriptomics data.

The SGE data were first normalised using the regularised negative binomial regression method implemented in the SCTransform function from Seurat (v3.1.4) R package. The data used for training of the model was the corrected UMI returned by the vst function from the sctransform package with the max and min values clipped to 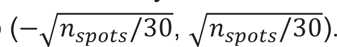

The DeepSpaCE model was then implemented using the VGG16 model as the backbone with the following hyper-parameters: maximum number of training epochs of 50 (with training stopping before 50 if there is no improvement within 5 epochs), a learning rate of 10^−4^ and a weight decay of 10^−4^.

The final model used was the model with the highest correlation in the validation set, as per the DeepSpaCE pipeline.

#### Hist2ST

Hist2ST is a deep-learning model predicting SGE from histology images comprising three main frameworks: Convmixer, Transformer and Graph neural network (GNN). It takes histology image patches, each sized 224x224, as inputs to the Convmixer framework, extracting 2D vision features from each patch. These features capture important local and global information within the patch. The embedded patch features are then fused with the corresponding embedded spot coordinates to create an enriched representation. These combined features are subsequently fed into the transformer framework, which considers the spatial relationships among the spots. Finally, the GNN module learns the relationships among neighbouring patches to enhance local spatial dependencies from the entire image. To improve model generalizability and performance, zero-inflated negative binomial distribution (ZINB) and self-distillation models were applied after the GNN. This allowed for capturing the probability of observing zero counts separately from the distribution of non-zero counts. Additionally, to mitigate the impact caused by small spatial transcriptomics data by fusing the features between the anchor image patch and the augmented image patches.

The model was implemented using Python and PyTorch and ran with default parameters stated in the original paper. These included: batch size of 1, learning rate of 0.00001, 350 epochs of training, and applying random grayscale, rotation and horizontal flip to image patches for self-distillation strategy. Some hyperparameter changes were made for benchmarking including: increased the image patch size from 112x112 to 224x224 and reduced the number of input and output channels in the depthwise and pointwise convolutional blocks from 32 to 16 due to limited computational power. Image embedding dimension was increased from 1024 to 2048 as the original code of setting embedding size was hard coded based on the image patch size and the number of channels. The model was trained using a sum of MSE loss of predicted and ground truth expression levels, ZINB and Distillation mechanisms.

#### ST-Net

ST-Net was one of the earliest methods proposed to infer spatial transcriptomics from H&E-stained histopathology images. ST-Net leveraged a CNN to capture gene expression heterogeneity in histology. A DenseNet-121 backbone was used to extract a 1,024-dim embedding of image patches (224 × 224 pixels in size), then a fully connected layer (with the same number of outputs as the number of genes) was applied to predict gene expressions of a spot. The weights of all convolutional layers were pre-trained on ImageNet. The authors originally used this architecture to predict the expressions of 250 genes.

ST-Net was implemented in Python and PyTorch using hyperparameter values as outlined in the original paper. These included: 224 × 224 pixel sized input image patches, batch size of 32, learning rate of 10e-6, 50 epochs of training, and applying image rotations and flips during training. The DenseNet-121 backbone used was pre-trained on ImageNet. ST-Net was trained using the MSE loss function, computed between predicted and ground truth expression levels.

#### DeepPT

The DeepPT architecture consisted of three main components in the following order: a CNN, auto- encoder, and a multi-layer perceptron (MLP). The CNN (ResNet-50 that was pre-trained on ImageNet) was used to extract features from image tiles, and produced a 2,048-dim embedding. The autoencoder comprised an input, hidden, and output layer that compressed the image embedding into a 512-dim vector. This step was designed to reduce data sparsity, avoid overfitting, and reduce noise. The compressed vector was then fed into a 3-layer MLP (consisting of an input, hidden, and output layer) to predict SGE. The authors treated each gene as an individual task in a multi-task paradigm; hence DeepPT is able to learn shared features between different genes.

DeepPT was implemented in Python and PyTorch using hyperparameter values as outlined in the original paper. These included: training for a maximum of 500 epochs, learning rate of 10e-4, batch size of 32, dropout set to 0.2, and applying rotations to the input images during training as data augmentation. The size of input image patches was 224 × 224 pixels. DeepPT was trained using the MSE loss function, computed between predicted and ground truth expression levels.

### Performance evaluation

<u>Overall rankings</u> were calculated by first taking the ranks of each method in each metric. These ranks were then averaged over all metrics within the same evaluation category to obtain an overall score for each category. Then the overall rankings of each category were then averaged over all categories to obtain a final overall score/rank.

Figures 2a and 2b, were created using the R package *funkyheatmap* ^30^. Scales of the bars in Figure 2a are relative within each category where the maximum value for x-axis for the bars represent the lowest average rank (with rank 1 being the best), and the minimum value of the x-axis for the bars represent the highest average rank (not necessarily zero).

### Evaluation metrics

Scale independent metrics were chosen to compare performance from different methods since some models were trained on normalised/augmented counts rather than the raw gene counts, see Table 2. This was done opposed to back-transforming normalised counts as it may introduce potential issues, such as loss of information or bias. This was particularly important as methods transformed the original data for specific reasons.

### Gene expression prediction

The SGE prediction was evaluated using 4-fold cross validation (CV). The spatial transcriptomics datasets were split into four folds, with samples from the same patient belonging in the same fold. In each iteration of the CV, two folds were used as a training set, one fold used as a validation set and the remaining fold used as a test set. The training set was used to train parameters of the model. The validation set was used to evaluate models during training to allow parameters to be tuned. The best model within each fold was then chosen based on the epoch that produced the best Pearson Correlation Coefficient (PCC) in the validation set. Results presented in the paper were calculated using the predictions from the best performing model (using the test set) out of the four folds of the CV, unless specified otherwise) The same CV splits were used for each model to ensure fair comparison. Metrics under this category of evaluation were first calculated at an image level (e.g. correlation was measured for each gene per image), and then averaged over each patient, then averaged over each gene.

For the clustering in Figure 3a, the default clustering in the *Heatmap* function from the ComplexHeatmap R package ^54^ was used. More specifically, the heatmap utilised average-linkage hierarchical clustering over the euclidean distances, with the hierarchy reordered through the *reorder.dendrogram* function from the *stats* R package ^55^.

HVGs in Figure 3c were deduced from the ground truth SGE from the ST datasets. The *modelGeneVar* function, followed by the *getTopHVGs* function with prop=0.1 from the *scran* R package ^56^ were used to obtain the HVGs in each ST dataset.

### Model generalisability

For TCGA images, a pseudo-bulk expression for each image was calculated by predicting the SGE for each image, and then averaging the gene expression across each image tile. Pseudo-bulk values were then transformed as follows:

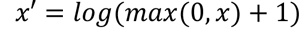

where x is the original predicted pseudobulk values for each gene and image.

The pseudo-bulk gene expression was then evaluated against the true bulk gene expression associated with the H&E image from the TCGA database. For the patient-level correlations, only the log-transformed gene values that were greater than 5 were included. This is appropriate as a way to filter out noise for a more informed evaluation.

Histology QC metrics were calculated using HistoQC python package. Whole slide images were converted to single-file pyramidal tiled TIFF format using nip2 interface for the libvips ^57^ package. Then HistoQC python package was run on each of the converted images for stage I breast cancer samples only. For Figure 4a, the first principal component from a PCA using colour-related metrics was plotted against the patient-level correlation between TCGA RNA-Seq and predicted pseudobulk GE. Colour-related metrics included: rms_contrast (the standard deviation of the pixel intensities across the pixels of interests), michelson_contrast (measurement of image contrast defined by luminance difference over average luminance), grayscale_brightness (mean pixel intensity of the image after converting the image to grayscale), chan1_brightness, chan2_brightness, chan3_brightness (mean pixel intensity of the image of the red, green and blue colour channels respectively), chan1_brightness_YUV, chan2_brightness_YUV and chan3_brightness_YUV (mean channel brightness of red, green and blue colour channels of image after converting to YUV colour space respectively)

### Clinical and/or translational impact

Out of the curated TCGA images, we split samples according to breast cancer (BC) subtypes as described in Xu et al ^29^. The clinical information such as ER-, PR-, HER2- were defined according to the estrogen, progesterone and HER2 receptor status variables and these were used to define BC subsets.

For each BC subset, we used the RNA-Seq to build a survival model using only the genes present in the HER2+ ST dataset. Alongside the model built from RNA-Seq, we used the predicted pseudobulk GE from each of the benchmarked methods to build additional survival models for comparison.

Pseudo-bulk values were normalised through mean centering (**Supplementary** Figure 8**, 9**). Days to death and days to last follow up (where there was no death) were used as the survival times, and death used as the survival outcome. We used the top five genes to build survival models within each BC subset and GE data. These top genes were determined by the c-indices of univariate cox models for each gene. A multivariate cox regression was then built using these top five genes.

Each model was assessed through the calculation of c-index and a logrank p-value for each BC subset and GE data (bulk RNA-Seq GE and predicted pseudobulk GE). C-Index was calculated through comparing the ground truth event times and events with model predictions for each patient. Predictions from the cox regression models were then used to categorise patients into high-risk and low-risk, using the median prediction to split patients. Kaplan-Meier curves were then constructed within high-risk and low-risk. The logrank test was then used to obtain a p-value to quantify the difference between survival of the two categories.

Two types of evaluation were used for survival models within each BC subset and GE data. We calculated c-indices and logrank p-values based on: (i) methods described in Xu et al. which is equivalent to a resubstitution model and (ii) a cross validation framework. For (i), models were built using all of the patients and then predictions for these patients were used for calculation of C-index and logrank p-value. For (ii), we used a 3-fold CV with 100 repeats. This meant that patients were randomly split into three subsets. Each subset was used as a test set of patients while the rest of the patients were used to train models. Through repeating this process 100 times, we measured the C-indices for each test set to obtain an average C-index over 300 values. We also obtained the average prediction for each patient over all repetitions which was then used to categorise patients into high and low-risk to calculate a logrank p-value.

### Usability

The user accessibility data was generated extending the scoring scheme used by Saelens et al. ^30^ to assess usability of trajectory inference methods. This scoring scheme is based on existing tool quality and programming guidelines found in literature. In addition to this scheme we added two sections: (i) reproducibility, assessing whether we were able to run the analysis pipeline with minimal changes to code; and (ii) generalisability, assessing whether methods included functionality to predict on and preprocess new images. These were important categories addressing whether users would be able to run these methods under a clinical translational setting. Additionally, we applied a slightly higher weight to these categories as part of scoring methods.

### Performance

For calculating performance, memory and time taken to train one image over 10 epochs was used as a unit for comparison across methods (**Supplementary** Figure 10). We achieved this by averaging the values across all epochs and then taking the ranks. Peak and instantaneous memory was calculated via the *get_traced_memory* from the *tracemalloc* module for all python methods whilst the *peakRAM* function from the *peakRAM* library was used for GeneCodeR, an R method.

All performance results reported in this section were obtained using Linux with Intel(R) Xeon(R) Gold 6338 CPU @ 2.00GHz and 1TB memory. Additionally, four NVIDIA RTX A5000 GPUs with 24GB of memory each were available for GPU-based methods.

All analysis of model predictions was performed using R ^55^ version 4.3.1. visualisations not already referenced were generated using ggplot2 ^58^.

### Code and data availability

Code and data for reproducing the results of this study is publicly available on Zenodo ^59^.

## Acknowledgement

The authors thank all their colleagues, particularly at the Sydney Precision Data Science Centre, Charles Perkins Centre and Biomedical Data Analysis and Visualisation Lab for their support and intellectual engagement. Special thanks to Yingxin Lin, Yue Cao, Lijia Yu, Andy Tran, Hao Wang, Andrew Sawyer and Yunwei Zhang for their contributions in weekly discussions.

## Authors’ contributions

JY and EP conceived and led the study with design input from SG and JK. AC led the development of the benchmarking framework input and guidance from JY and EP. AC, CW and XF shared the data curation and processing implementation of all existing methods and ran the benchmarking studies. AC and CW co-led the development and interpretation of the evaluation framework with input from JY, EP, SG, JK and XF. All authors contributed to the writing, editing, and approval of the manuscript.

## References

1. Ståhl, P. L. et al. Visualization and analysis of gene expression in tissue sections by spatial transcriptomics. Science 353, 78–82 (2016).

2. Chen, W.-T. et al. Spatial Transcriptomics and In Situ Sequencing to Study Alzheimer’s Disease. Cell 182, 976–991.e19 (2020).

3. Baccin, C. et al. Combined single-cell and spatial transcriptomics reveal the molecular, cellular and spatial bone marrow niche organization. Nat. Cell Biol. 22, 38–48 (2020).

4. Walker, B. L., Cang, Z., Ren, H., Bourgain-Chang, E. & Nie, Q. Deciphering tissue structure and function using spatial transcriptomics. Commun Biol 5, 220 (2022).

5. He, B. et al. Integrating spatial gene expression and breast tumour morphology via deep learning. Nat Biomed Eng 4, 827–834 (2020).

6. Pang, M., Su, K. & Li, M. Leveraging information in spatial transcriptomics to predict super-resolution gene expression from histology images in tumors. bioRxiv 2021.11.28.470212 (2021) doi:10.1101/2021.11.28.470212.

7. Hoang, D., Dinstag, G., Hermida, L. C., Ben-Zvi, D. S. & Elis, E. Synthetic lethality-based prediction of cancer treatment response from histopathology images. https://europepmc.org/article/ppr/ppr504688.

8. Zeng, Y. et al. Spatial transcriptomics prediction from histology jointly through Transformer and graph neural networks. Brief. Bioinform. 23, (2022).

9. Monjo, T., Koido, M., Nagasawa, S., Suzuki, Y. & Kamatani, Y. Efficient prediction of a spatial transcriptomics profile better characterizes breast cancer tissue sections without costly experimentation. Sci. Rep. 12, 4133 (2022).

10. 10. Banh, D. & Huang, A. Scalable parametric encoding of multiple modalities. bioRxiv 2021.07.09.451779 (2022) doi:10.1101/2021.07.09.451779.

11. Yang, Y., Hossain, M. Z., Stone, E. A. & Rahman, S. Exemplar guided deep neural network for spatial transcriptomics analysis of gene expression prediction. in 2023 IEEE/CVF Winter Conference on Applications of Computer Vision (WACV) 5039–5048 (IEEE, 2023).

12. Yang, Y., Hossain, M. Z., Stone, E. & Rahman, S. Spatial Transcriptomics Analysis of Gene Expression Prediction using Exemplar Guided Graph Neural Network. bioRxiv 2023.03.30.534914 (2023) doi:10.1101/2023.03.30.534914.

13. Bergenstråhle, L. et al. Super-resolved spatial transcriptomics by deep data fusion. Nat. Biotechnol. 40, 476–479 (2022).

14. Xie, R., Pang, K., Bader, G. D. & Wang, B. Spatially Resolved Gene Expression Prediction from H&E Histology Images via Bi-modal Contrastive Learning. arXiv [cs.CV] (2023).

15. Dawood, M., Branson, K., Rajpoot, N. M. & Minhas, F. ul A. A. All You Need is Color: Image Based Spatial Gene Expression Prediction Using Neural Stain Learning. in Machine Learning and Principles and Practice of Knowledge Discovery in Databases 437–450 (Springer International Publishing, 2021).

16. Xiao, X., Kong, Y., Wang, Z. & Lu, H. Transformer with Convolution and Graph-Node co-embedding: An accurate and interpretable vision backbone for predicting gene expressions from local histopathological image. *bioRxiv* 2023.05.28.542669 (2023) doi:10.1101/2023.05.28.542669.

17. Rahaman, M. M., Millar, E. K. A. & Meijering, E. Breast Cancer Histopathology Image based Gene Expression Prediction using Spatial Transcriptomics data and Deep Learning. arXiv [eess.IV] (2023).

18. Zhao, C. et al. Transformer Enables Reference Free And Unsupervised Analysis of Spatial Transcriptomics. *bioRxiv* 2022.08.11.503261 (2022) doi:10.1101/2022.08.11.503261.

19. Pang, M., Su, K. & Li, M. Leveraging information in spatial transcriptomics to predict super-resolution gene expression from histology images in tumors. bioRxiv 2021.11.28.470212 (2021) doi:10.1101/2021.11.28.470212.

20. Moses, L. & Pachter, L. Museum of spatial transcriptomics. Nat. Methods 19, 534–546 (2022).

21. Li, Y., Stanojevic, S. & Garmire, L. X. Emerging artificial intelligence applications in Spatial Transcriptomics analysis. Comput. Struct. Biotechnol. J. 20, 2895–2908 (2022).

22. Nasab, R. Z., et al. Deep Learning in Spatially Resolved Transcriptomics: A Comprehensive Technical View. *arXiv [q-bio.GN]* (2022).

23. 23. Hoang, D.-T., et al. Prediction of cancer treatment response from histopathology images through imputed transcriptomics. bioRxiv 2022.06.07.495219 (2023) doi:10.1101/2022.06.07.495219.

24. Schroeder, B. et al. Fatty acid synthase (FASN) regulates the mitochondrial priming of cancer cells. Cell Death Dis. 12, 977 (2021).

25. Gurda, D., Handschuh, L., Kotkowiak, W. & Jakubowski, H. Homocysteine thiolactone and N-homocysteinylated protein induce pro-atherogenic changes in gene expression in human vascular endothelial cells. Amino Acids 47, 1319–1339 (2015).

26. Dang, C. et al. Identification of dysregulated genes in cutaneous squamous cell carcinoma. Oncol. Rep. 16, 513–519 (2006).

27. Janowczyk, A., Zuo, R., Gilmore, H., Feldman, M. & Madabhushi, A. HistoQC: An Open-Source Quality Control Tool for Digital Pathology Slides. JCO Clin Cancer Inform 3, 1–7 (2019).

28. Bulten, W. et al. Epithelium segmentation using deep learning in H&E-stained prostate specimens with immunohistochemistry as reference standard. Sci. Rep. 9, 864 (2019).

29. Xu, J. et al. Delving into the Heterogeneity of Different Breast Cancer Subtypes and the Prognostic Models Utilizing scRNA-Seq and Bulk RNA-Seq. Int. J. Mol. Sci. 23, (2022).

30. Saelens, W., Cannoodt, R., Todorov, H. & Saeys, Y. A comparison of single-cell trajectory inference methods. Nat. Biotechnol. 37, 547–554 (2019).

31. Vaswani, A. et al. Attention is All you Need. in Advances in Neural Information Processing Systems (eds. Guyon, I. et al.) vol. 30 (Curran Associates, Inc., 2017).

32. Minoura, K., Abe, K., Nam, H., Nishikawa, H. & Shimamura, T. A mixture-of-experts deep generative model for integrated analysis of single-cell multiomics data. Cell Rep Methods 1, 100071 (2021).

33. Li, J., Chen, S., Pan, X., Yuan, Y. & Shen, H.-B. Cell clustering for spatial transcriptomics data with graph neural networks. Nature Computational Science 2, 399–408 (2022).

34. Kirillov, A., et al. Segment Anything. arXiv [cs.CV] (2023).

35. Ince, D. C., Hatton, L. & Graham-Cumming, J. The case for open computer programs. Nature 482, 485–488 (2012).

36. McKiernan, E. C. et al. How open science helps researchers succeed. Elife 5, (2016).

37. Heil, B. J. et al. Reproducibility standards for machine learning in the life sciences. Nat. Methods 18, 1132–1135 (2021).

38. Gustavsson, T. Managing the open source dependency. Computer 53, 83–87 (2020).

39. Wilson, G. et al. Good enough practices in scientific computing. PLoS Comput. Biol. 13, e1005510 (2017).

40. Ivie, P. & Thain, D. Reproducibility in Scientific Computing. ACM Comput. Surv. 51, 1– 36 (2018).

41. Sandve, G. K., Nekrutenko, A., Taylor, J. & Hovig, E. Ten simple rules for reproducible computational research. PLoS Comput. Biol. 9, e1003285 (2013).

42. Tan, X. et al. STimage:robust, confident and interpretable models for predicting gene markers from cancer histopathological images. bioRxiv 2023.05.14.540710 (2023) doi:10.1101/2023.05.14.540710.

43. Xun, Z. et al. Reconstruction of the tumor spatial microenvironment along the malignant-boundary-nonmalignant axis. Nat. Commun. 14, 933 (2023).

44. He, S. et al. Starfysh reveals heterogeneous spatial dynamics in the breast tumor microenvironment. bioRxiv 2022.11.21.517420 (2022) doi:10.1101/2022.11.21.517420.

45. 45. Comiter, C., et al. Inference of single cell profiles from histology stains with the Single-Cell omics from Histology Analysis Framework (SCHAF). bioRxiv (2023) doi:10.1101/2023.03.21.533680.

46. Hou, S., et al. St2cell: Reconstruction of in situ single-cell spatial transcriptomics by integrating high-resolution histological image. *bioRxiv* 2022.10.13.512059 (2022) doi:10.1101/2022.10.13.512059.

47. Na, K. J., Koh, J., Choi, H. & Kim, Y. T. Mapping cell types in the tumor microenvironment from tissue images via deep learning trained by spatial transcriptomics of lung adenocarcinoma. bioRxiv 2023.03.04.531083 (2023) doi:10.1101/2023.03.04.531083.

48. Schmauch, B. et al. A deep learning model to predict RNA-Seq expression of tumours from whole slide images. Nat. Commun. 11, 1–15 (2020).

49. Yang, Y., Pan, L., Liu, L. & Stone, E. A. ISG: I can See Your Gene Expression. arXiv [cs.CV] (2022).

50. Andersson, A. et al. Spatial deconvolution of HER2-positive breast cancer delineates tumor-associated cell type interactions. Nat. Commun. 12, 6012 (2021).

51. Ji, A. L. et al. Multimodal Analysis of Composition and Spatial Architecture in Human Squamous Cell Carcinoma. Cell 182, 1661–1662 (2020).

52. 52. Cancer Genome Atlas Network. Comprehensive molecular portraits of human breast tumours. Nature 490, 61–70 (2012).

53. Colaprico, A. et al. TCGAbiolinks: an R/Bioconductor package for integrative analysis of TCGA data. Nucleic Acids Res. 44, e71 (2016).

54. Gu, Z., Eils, R. & Schlesner, M. Complex heatmaps reveal patterns and correlations in multidimensional genomic data. Bioinformatics 32, 2847–2849 (2016).

55. 55. R Core Team. R: A Language and Environment for Statistical Computing. Preprint at https://www.R-project.org/ (2020).

56. Lun, A. T. L., McCarthy, D. J. & Marioni, J. C. A step-by-step workflow for low-level analysis of single-cell RNA-seq data with Bioconductor. F1000Res. 5, 2122 (2016).

57. Martinez, K. & Cupitt, J. LibVIPS: A fast image processing library with low memory needs. (2007).

58. Wickham, H. Data Analysis. in ggplot2: Elegant Graphics for Data Analysis (ed. Wickham, H.) 189–201 (Springer International Publishing, 2016).

59. Chan, Adam. Benchmarking the translational potential of spatial gene expression prediction from histology analysis. (Zenodo, 2023). doi:10.5281/zenodo.10213326.

60. Monjo, T., Koido, M., Nagasawa, S., Suzuki, Y. & Kamatani, Y. Efficient prediction of a spatial transcriptomics profile better characterizes breast cancer tissue sections without costly experimentation. Sci. Rep. 12, 4133 (2022).

