## Supplemental Tables and Figures for "Benchmarking the translational potential of spatial gene expression prediction from histology"

### Supplementary Figures and Tables

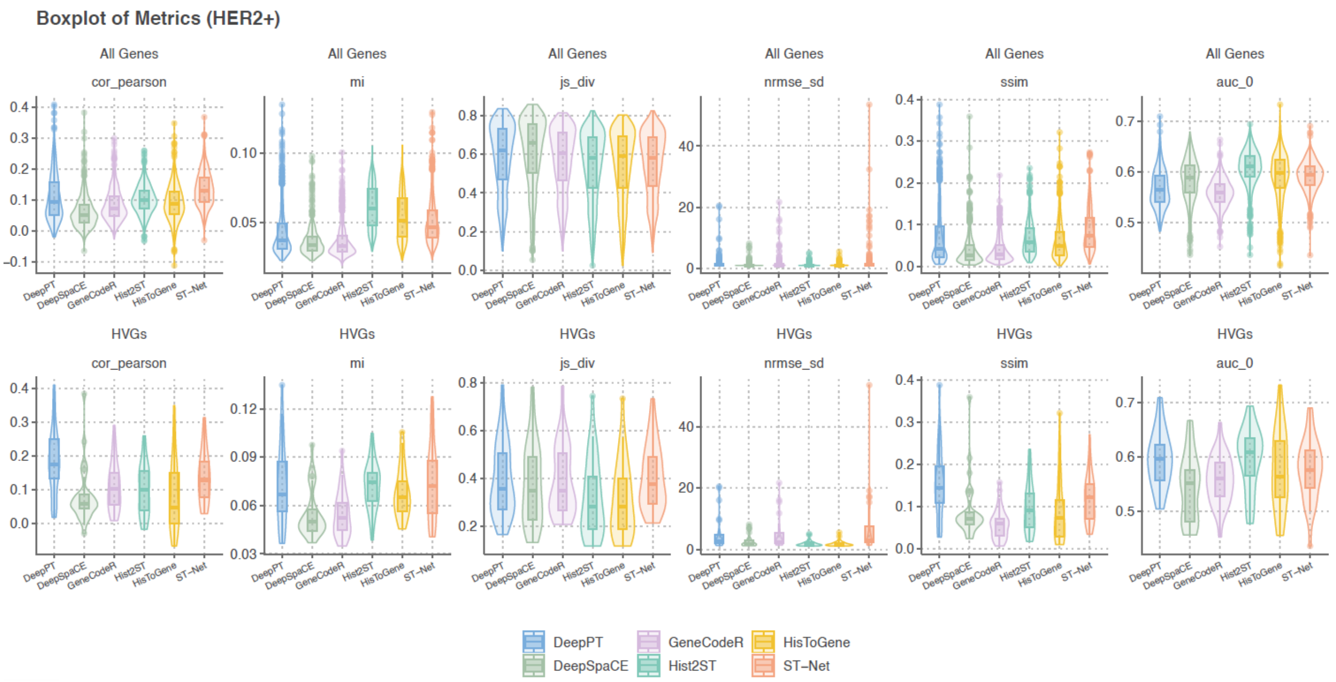


**Supplementary Figure 1:** Violin and boxplots of evaluation metrics for gene expression for each method in the HER2+ ST dataset

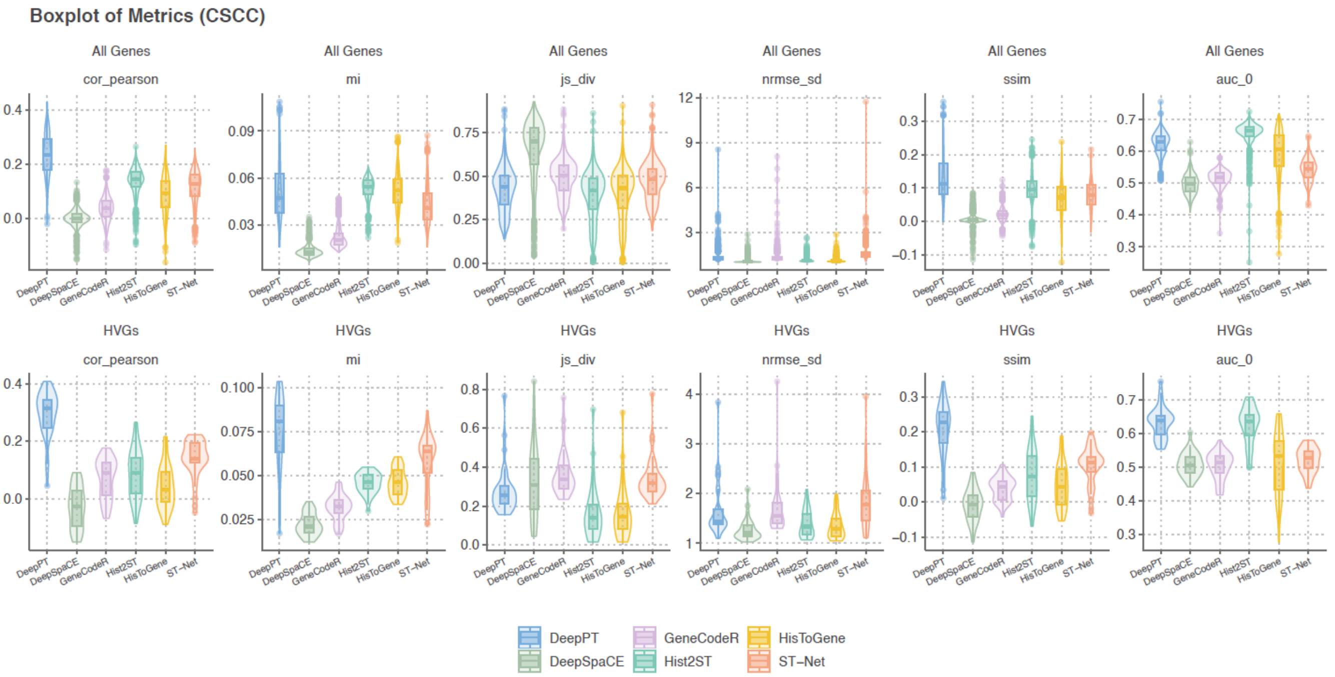


**Supplementary Figure 2:** Violin and boxplots of evaluation metrics for gene expression for each method in the CSCC ST dataset

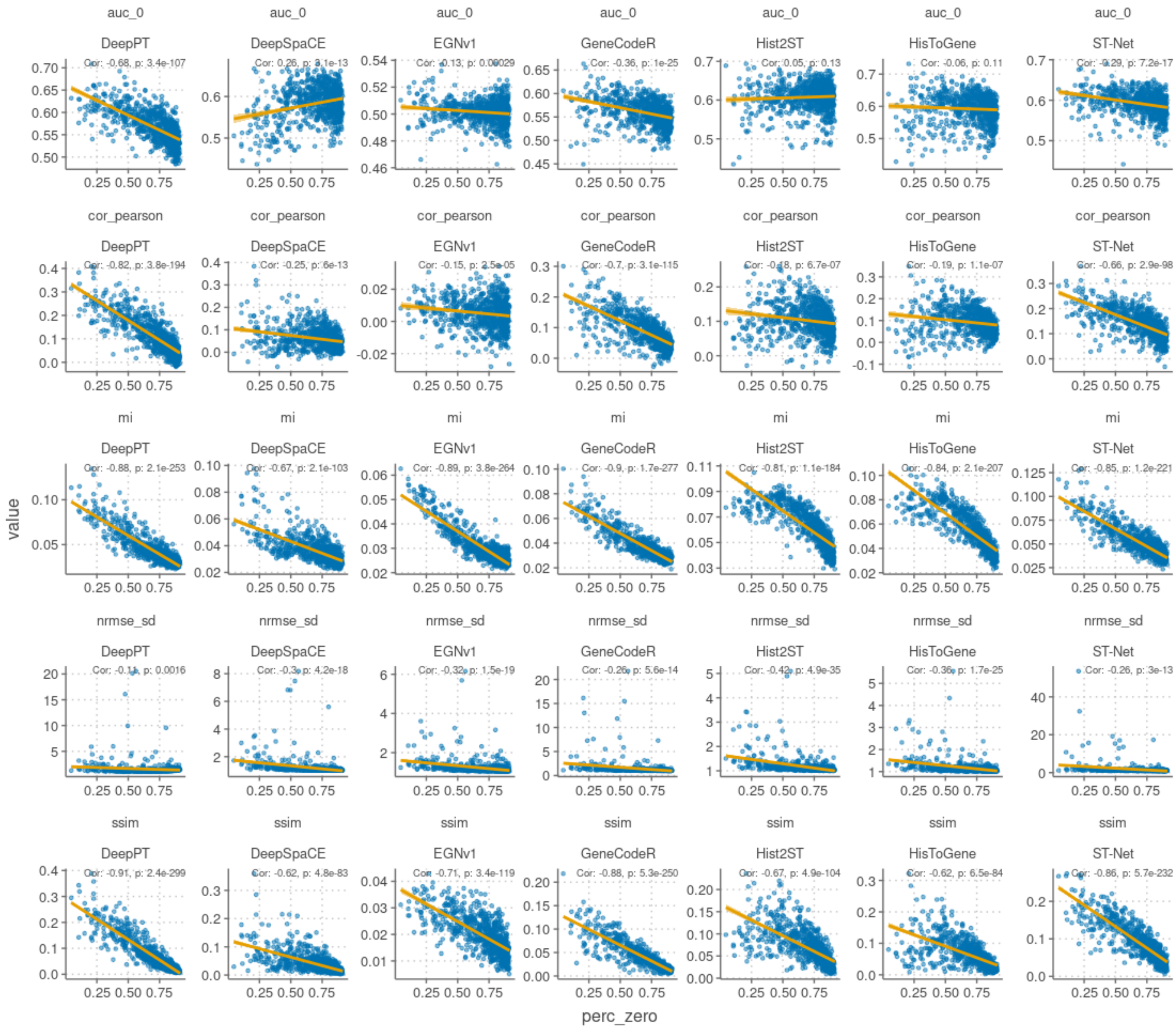


**Supplementary Figure 3:** Gene expression prediction evaluation metrics vs. the percentage of zeros in each gene for each method. Linear lines of best fit are plotted for each.

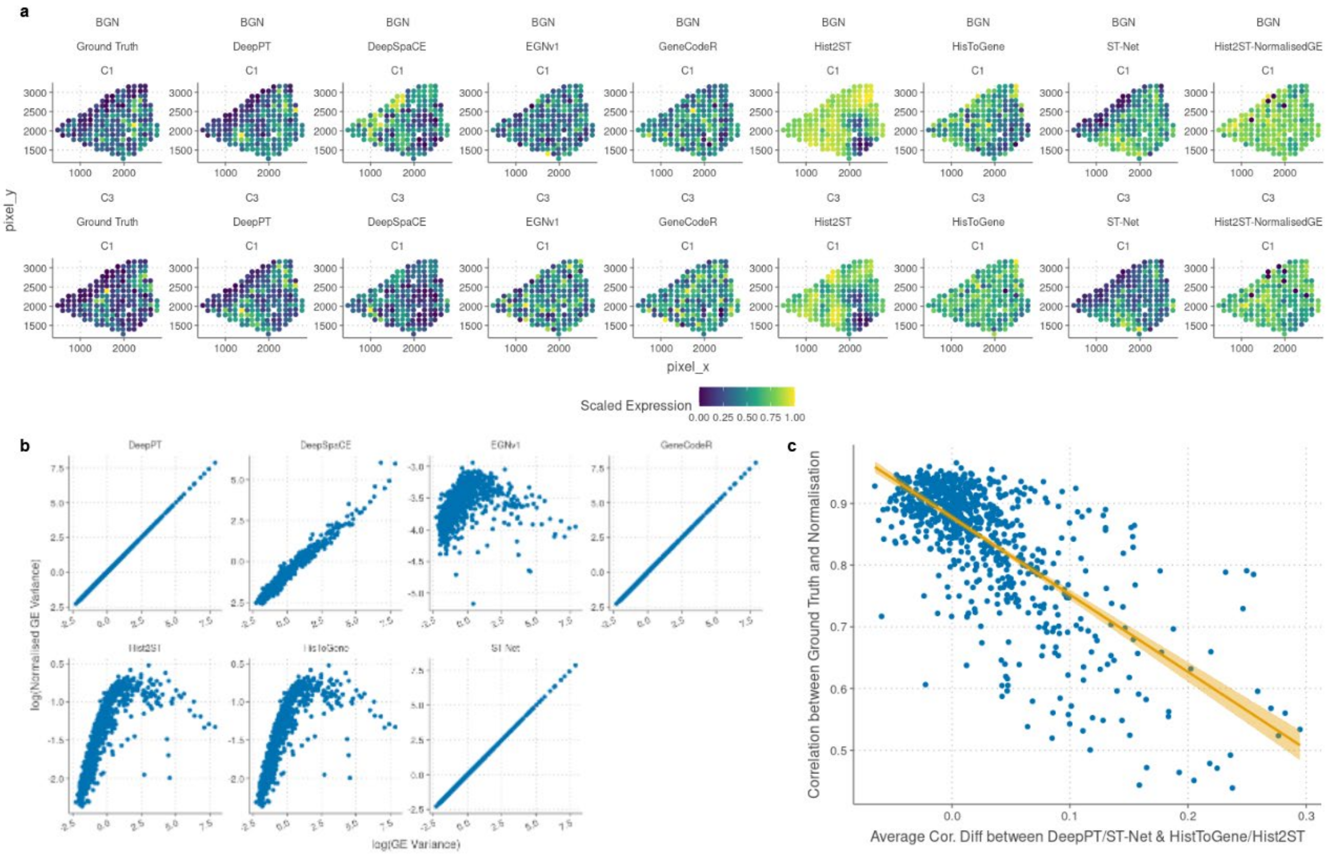


**Supplementary Figure 4:** Illustrating difference in correlation between Hist2ST/HisToGene & DeepPT/ST-Net in the HER2+ ST dataset. (a) Scatterplot of the gene expression at each spot for each of the model predictions in the training set for one image and genes C3 and BGN. Genes were chosen as they had high correlation in DeepPT/ST-Net and low correlation in Hist2ST/HisToGene. Ground truth (leftmost column) and normalised gene expression (rightmost column) values are also plotted. (b) Scatterplot of gene expression variance before (x-axis) and after normalisation (y-axis) for each method. (c) Scatterplot of average correlation difference between average correlation of both DeepPT/ST-Net & average correlation of both HisToGene/Hist2ST and the correlation between ground truth and normalisation (y-axis). Each point represents a gene.

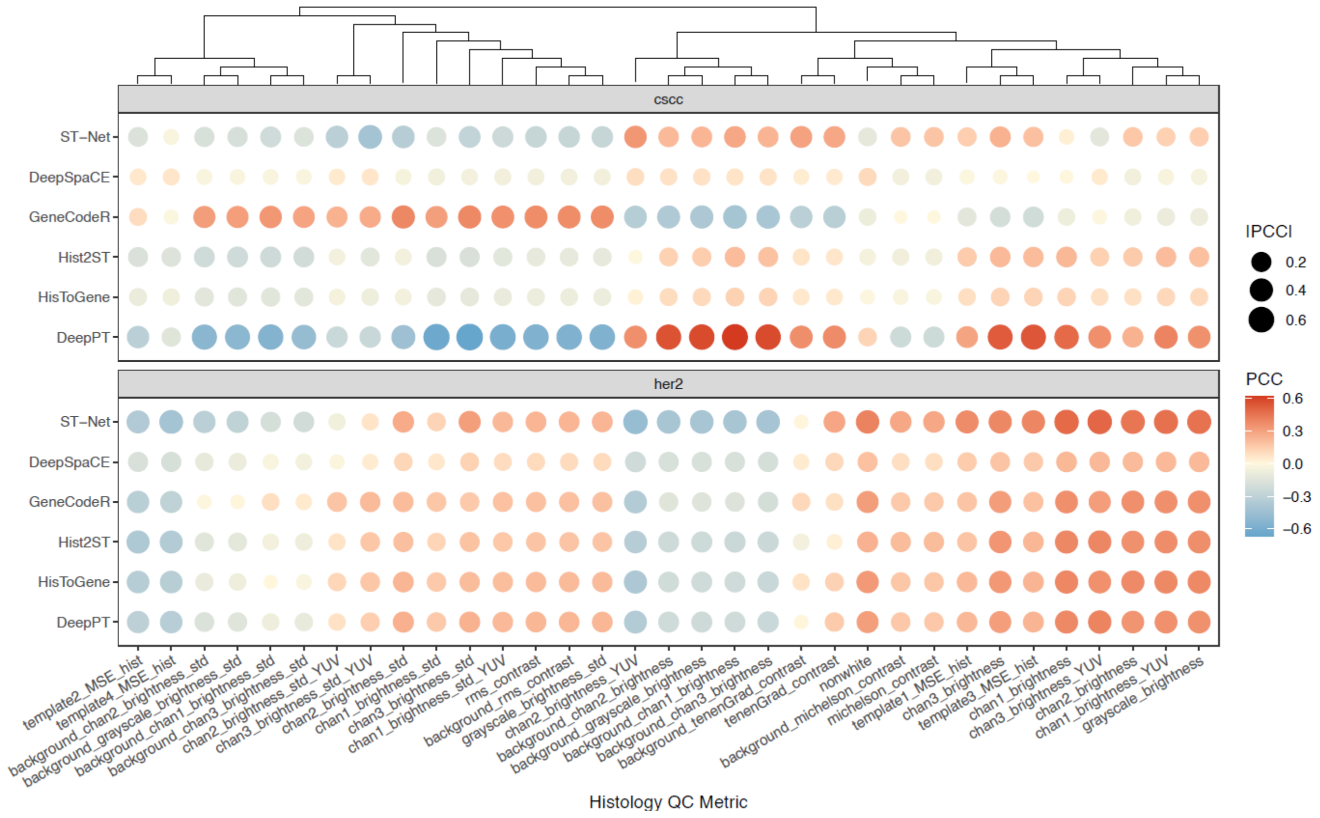


**Supplementary Figure 5:** Dotplot of correlation between various histology QC metrics and gene-level correlations for each method in the HER2+ ST dataset and the CSCC ST dataset.

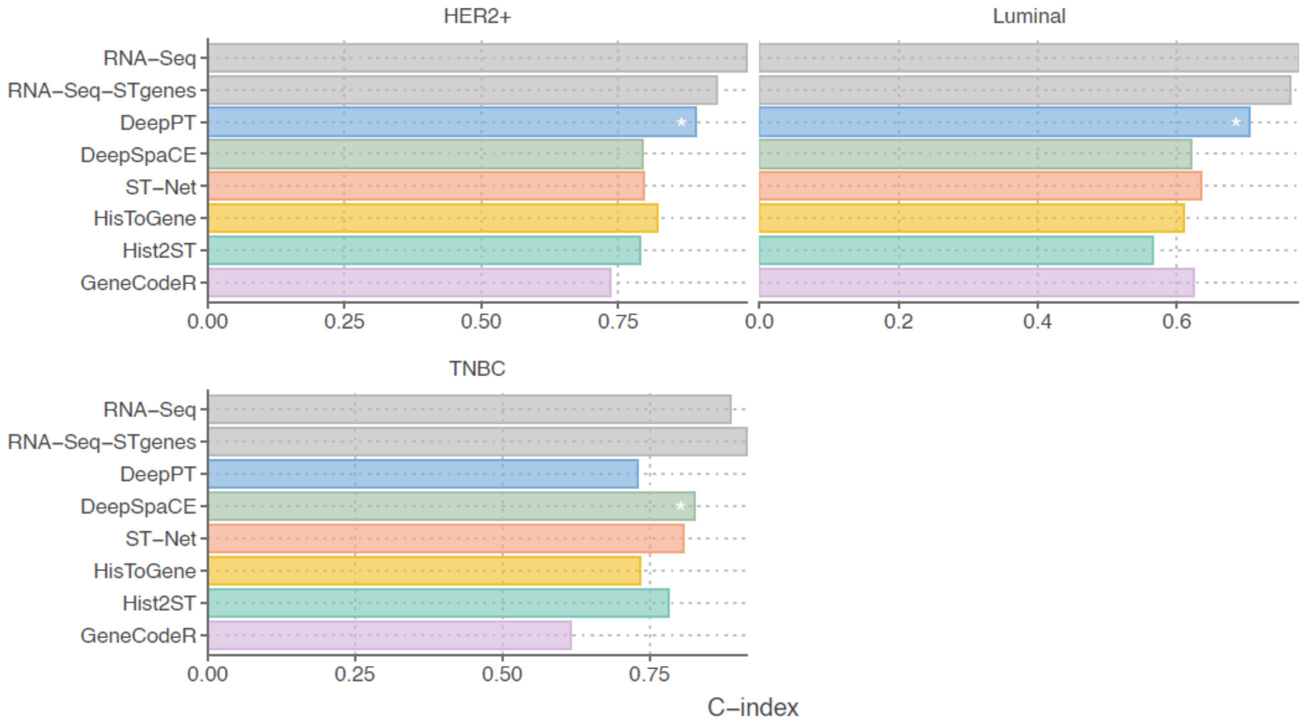


**Supplementary Figure 6:** C-indices of multivariate cox regression models predicting survival of TCGA-BRCA patients, using RNA-Seq bulk, RNA-Seq bulk using only genes present in HER2+ ST dataset, and the predicted pseudobulk from each method. C-indices were calculated from the training data of models trained within (c) HER2+, (d) luminal and (e) TNBC breast cancer clinical subtypes

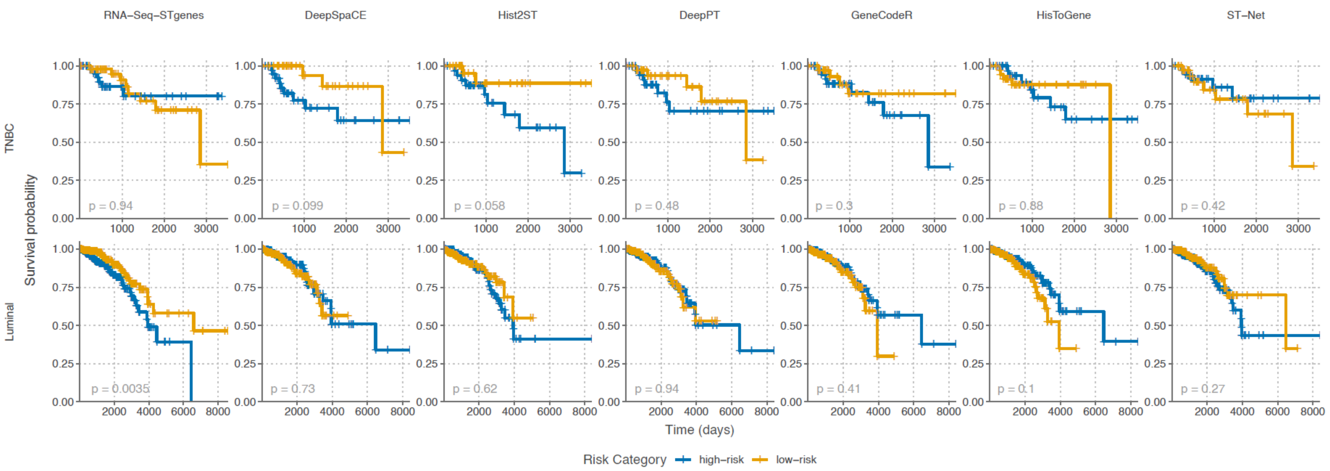


**Supplementary Figure 7:** Kaplan-Meier curves for patients split into high and low risk groups by the median risk prediction of the multivariate cox regression models for each method in luminal and TNBC breast cancer subtypes. The average risk prediction from a 3-fold CV with 100 repeats was used. The p-value represents the result of the logrank test for assessing the statistical significance of differences in survival between the groups.

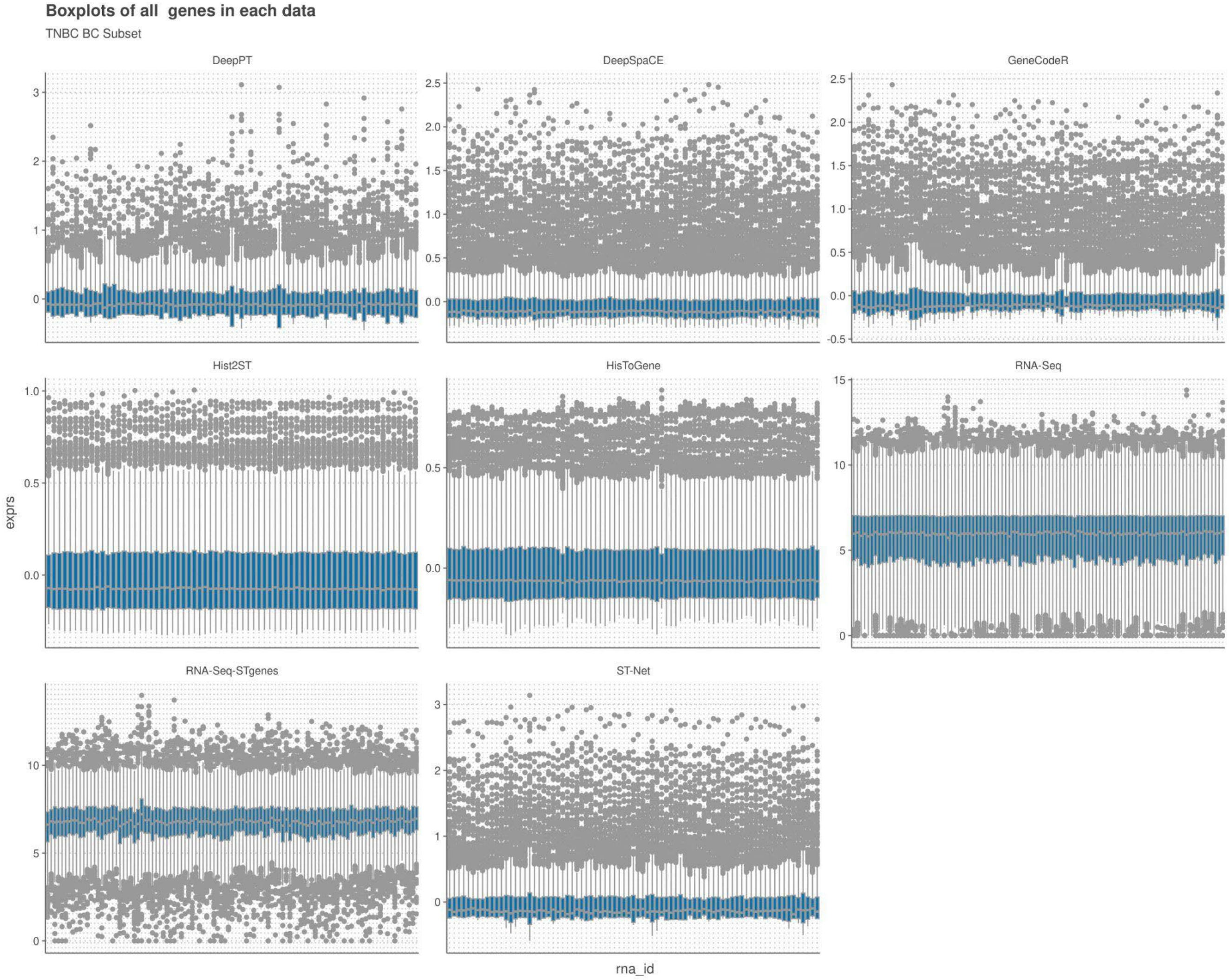


**Supplementary Figure 8:** Boxplot of gene expression values after transformation for each sample from the TNBC subset of the TCGA data and for all genes that were measured in the HER2+ spatial transcriptomics dataset.

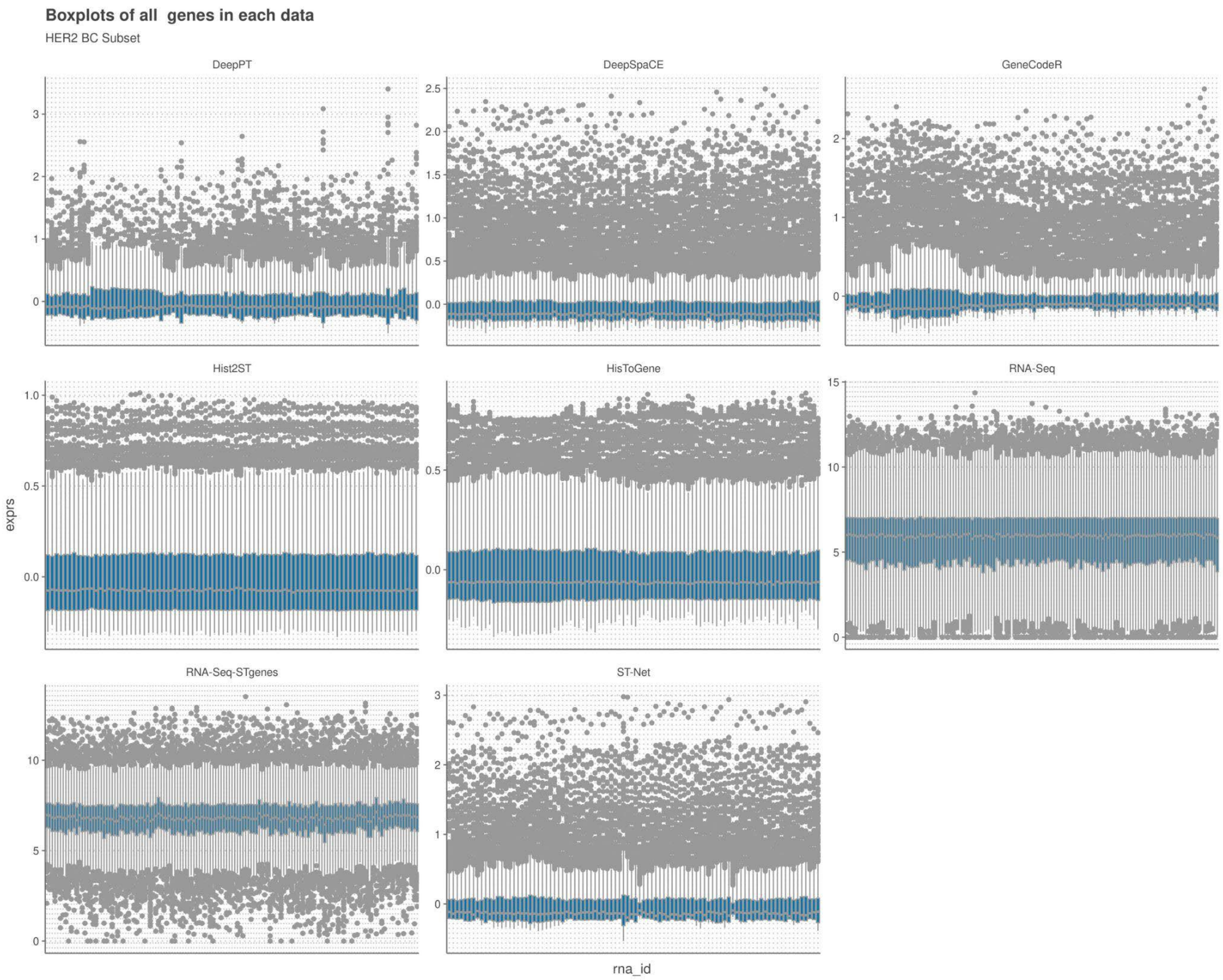


**Supplementary Figure 9:** Boxplot of gene expression values after transformation for each sample from the HER2 subset of the TCGA data and for all genes that were measured in the HER2+ spatial transcriptomics dataset.

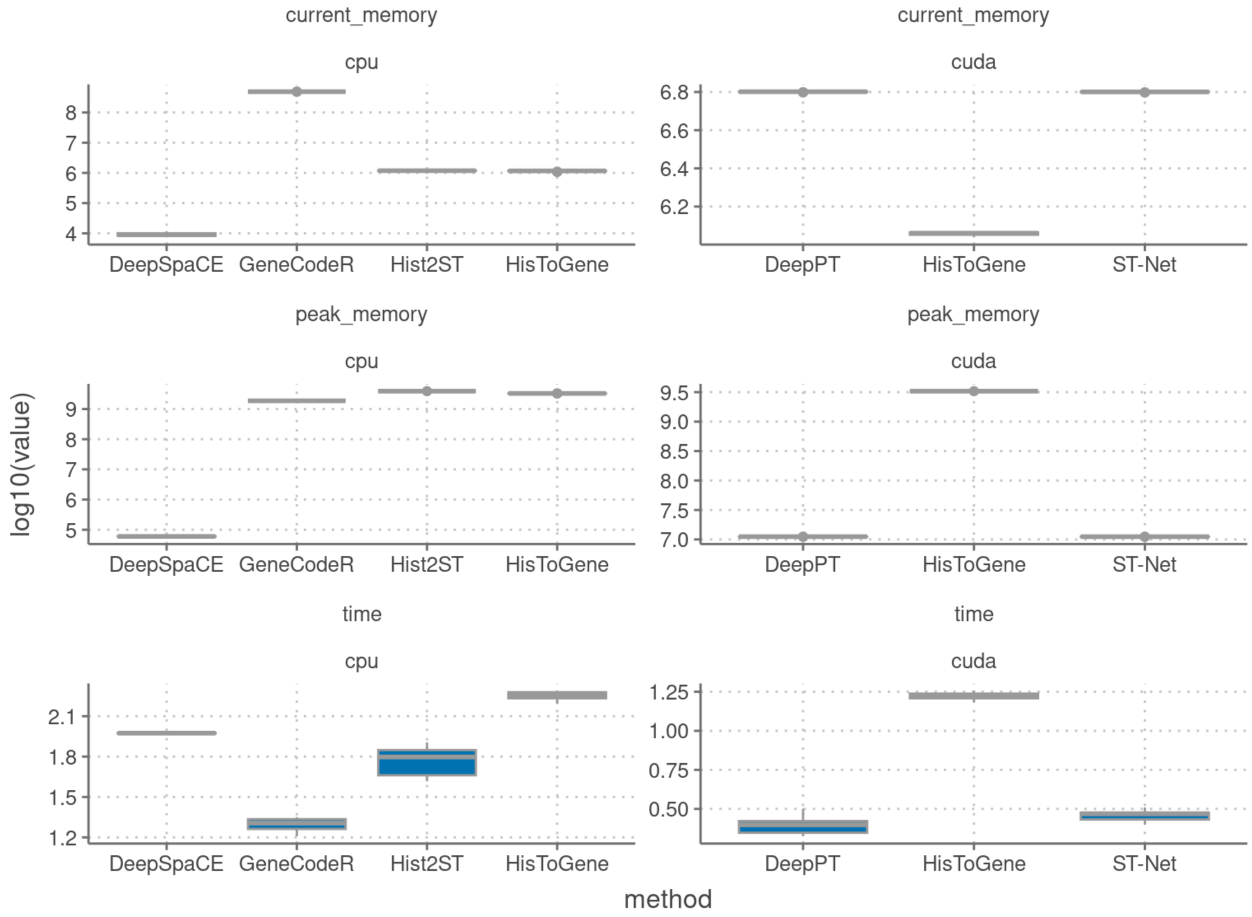


**Supplementary Figure 10:** Boxplots of the computational efficiency of methods when trained on one histology image using 10 epochs. Metrics were measured for parallelised (cuda) and non-parallelised (cpu) training where applicable. Memory was measured in bytes and time in seconds.

#### Supplementary Table 1: Summary of Methods predicting SGE from H&E

| Model | Local Features (one spot) | Local + Global Features (spot-neighborhood relations) | Global Features (spot-spatial relations) | Reference Dataset | Reference Encoder | Application method |
|---|---|---|---|---|---|---|
| ST-Net [5] | Pretrained DenseNet 121 | NA | NA | NA | NA | NA |
| HisToGene [19] | ViT | Super Resolution | NA | NA | NA | NA |
| DeepPT [7] | Pretrained ResNet50 + Autoencoder + MLP | NA | NA | NA | NA | NA |
| Hist2ST [8] | Convmixer | GNN - GraphSAGE | Transformer | NA | NA | NA |
| DeepSpaCE [60] | VGG16 | Super Resolution | NA | NA | NA | NA |
| GeneCodeR [10] | Non deep learning method | | | | | |
| EGN [11] | ViT + Exemplar | NA | Exemplar | NA | NA | NA |
| EGN v2[12] | Exemplar(ResNet) + GraphSAGE + GCN | NA | Exemplar | Reference dataset | NA | Exemplar Retrieval |
| XFuse [13] | Statistical Model + Deep generative model | | | ISC Data | NA | Fuse with H&E |
| BLEEP [14] | Pretrained ResNet50 | Contrastive Learning | | NA | NA | NA |
| NSL [15] | Stain deconvolution matrix | | | | | |
| TCGN [16] | CNN + ViT + GNN | NA | NA | NA | NA | NA |

| | | | | | | |
|---|---|---|---|---|---|---|
| BrST-Ne [17] | Trained 10 state-of-the-art CNN models and transformers then compared their performances + introduced an auxiliary network | | | | | |
| TransformerST [18] | CNN + cross-scale internal GNN | Adaptive graph transformer | Conditional transformer | NA | NA | NA |
| STimage [42] | Pretrained ResNet50 + Negative Binomial | NA | NA | NA | NA | NA |

#### Supplementary Table 2: Top 20 predicted genes by correlation in HER2+ data.

| gene | DeepPT | DeepSpaCE | GeneCodeR | HisToGene | Hist2ST | ST-Net | overall_mean_cor |
|---|---|---|---|---|---|---|---|
| GNAS | **0.41** | 0.38 | 0.29 | 0.35 | 0.26 | 0.28 | 0.33 |
| FASN | **0.4** | 0.32 | 0.28 | 0.31 | 0.21 | 0.37 | 0.32 |
| SCD | **0.34** | 0.24 | 0.15 | 0.28 | 0.22 | 0.31 | 0.26 |
| MYL12B | **0.3** | 0.23 | 0.19 | 0.28 | 0.26 | 0.27 | 0.25 |
| CLDN4 | **0.31** | 0.2 | 0.23 | 0.28 | 0.21 | 0.27 | 0.25 |
| RHOB | **0.29** | 0.25 | 0.19 | 0.25 | 0.25 | 0.21 | 0.24 |
| STMN1 | **0.28** | 0.25 | 0.15 | 0.26 | 0.23 | 0.22 | 0.23 |
| FN1 | 0.26 | 0.18 | 0.17 | **0.27** | 0.24 | 0.2 | 0.22 |
| HLA.DRA | **0.36** | 0.13 | 0.21 | 0.19 | 0.18 | 0.23 | 0.22 |
| NDUFB2 | **0.26** | 0.14 | 0.18 | 0.24 | 0.25 | 0.22 | 0.22 |
| CCT4 | **0.27** | 0.19 | 0.14 | 0.25 | 0.22 | 0.23 | 0.22 |
| PRKCSH | **0.3** | 0.11 | 0.24 | 0.21 | 0.16 | 0.26 | 0.21 |
| TMBIM6 | **0.28** | 0.14 | 0.17 | 0.23 | 0.2 | 0.25 | 0.21 |
| HMGB2 | **0.24** | 0.23 | 0.13 | 0.23 | 0.23 | 0.21 | 0.21 |

| | | | | | | | |
|---|---|---|---|---|---|---|---|
| HNRNPUL2 | **0.27** | 0.17 | 0.2 | 0.21 | 0.18 | 0.22 | 0.21 |
| SRSF1 | **0.26** | 0.2 | 0.14 | 0.23 | 0.21 | 0.23 | 0.21 |
| FADS2 | **0.27** | 0.1 | 0.26 | 0.21 | 0.15 | 0.25 | 0.21 |
| TXNDC17 | **0.23** | 0.19 | 0.16 | **0.23** | 0.21 | 0.22 | 0.21 |
| SRSF5 | **0.25** | 0.12 | 0.21 | 0.22 | 0.2 | 0.23 | 0.21 |
| CRACR2B | **0.26** | 0.14 | 0.2 | 0.21 | 0.16 | **0.26** | 0.2 |

#### Supplementary Table 3: Top 20 predicted genes by correlation in CSCC data.

| gene | DeepPT | DeepSpaCE | GeneCodeR | HisToGene | Hist2ST | ST-Net | overall_mean_cor |
|---|---|---|---|---|---|---|---|
| PFN1 | **0.41** | 0.08 | 0.12 | 0.12 | 0.27 | 0.13 | 0.19 |
| TAGLN2 | **0.38** | 0.04 | 0.1 | 0.23 | 0.22 | 0.16 | 0.19 |
| RPL24 | **0.37** | 0.03 | 0.06 | 0.18 | 0.21 | 0.22 | 0.18 |
| RPS17 | **0.34** | 0.05 | 0.1 | 0.19 | 0.25 | 0.13 | 0.18 |
| RPL9 | **0.36** | 0.06 | 0.07 | 0.16 | 0.24 | 0.16 | 0.17 |
| MYL12B | **0.35** | 0.01 | 0.06 | 0.22 | 0.2 | 0.21 | 0.17 |
| PRDX1 | **0.34** | 0.01 | 0.06 | 0.22 | 0.21 | 0.19 | 0.17 |
| ANXA2 | **0.39** | 0 | 0.12 | 0.15 | 0.18 | 0.19 | 0.17 |
| LMNA | **0.38** | 0.03 | 0.12 | 0.17 | 0.14 | 0.2 | 0.17 |
| RPS4X | **0.36** | 0.06 | 0.01 | 0.22 | 0.22 | 0.17 | 0.17 |
| PPIA | **0.34** | 0.01 | 0.12 | 0.23 | 0.22 | 0.12 | 0.17 |

| | | | | | | | |
|---|---|---|---|---|---|---|---|
| PKP1 | **0.33** | -0.09 | 0.13 | 0.27 | 0.19 | 0.19 | 0.17 |
| RPN2 | **0.35** | 0.04 | 0.07 | 0.2 | 0.19 | 0.19 | 0.17 |
| ACTG1 | **0.41** | 0 | 0.08 | 0.14 | 0.2 | 0.2 | 0.17 |
| OAZ1 | **0.34** | 0.03 | 0.05 | 0.19 | 0.22 | 0.18 | 0.17 |
| RPL5 | **0.34** | 0.06 | 0.08 | 0.17 | 0.23 | 0.14 | 0.17 |
| MYL6 | **0.38** | 0.02 | 0.05 | 0.19 | 0.2 | 0.17 | 0.17 |
| RPL18 | **0.33** | 0.01 | 0.01 | 0.19 | 0.24 | 0.23 | 0.17 |
| S100A16 | **0.35** | -0.03 | 0.09 | 0.18 | 0.22 | 0.2 | 0.17 |
| PTMA | **0.36** | 0.03 | 0.11 | 0.14 | 0.24 | 0.11 | 0.17 |

**Supplementary Table 4:** Usability scoring schema used to evaluate methods

| **name** | **aspect_id** | **category** | **aspect** | **item_weight** | **item** | **GeneCodeR** | **ST-Net** | **DeepPT** | **Hist2ST** | **HisToGene** | **DeepSpaCE** |
|---|---|---|---|---|---|---|---|---|---|---|---|
| Open source | open_source | availability | Is the code freely available? | 0.5 | Method's code is freely available | 1 | 1 | 0 | 1 | 1 | 1 |
| | | | | 0.5 | The code can be run on a freely available platform | 1 | 1 | 0 | 1 | 1 | 1 |
| Version control | version_control | availability | Is the code version controlled? | 1 | The code is available on a public version controlled repository, such as Github | 1 | 1 | 0 | 1 | 1 | 1 |
| Packaging | packaging | availability | Is the code contained into an easy to install package? Is it discoverable? | 0.5 | The code is provided as a "package", exposing functionality through functions or shell commands | 1 | 1 | 0 | 1 | 1 | 1 |
| | | | | 0.5 | The code can be easily installed through a repository such as CRAN, Bioconductor, PyPI, CPAN, debian packages, ... | 1 | 0 | 0 | 0 | 0 | 0 |
| Dependencies | dependencie | availability | Are dependencies clearly stated and | 0.5 | Dependencies are clearly stated in the tutorial or in the code | 0 | 0 | 0 | 1 | 1 | 1 |

| | | | | | | | | | | | |
|---|---|---|---|---|---|---|---|---|---|---|---|
| | s | | available? | 0.5 | Dependencies are automatically installed | 1 | 0 | 0 | 0 | 0 | 1 |
| License | license | availability | Is the license of the code clear? Does this license permit academic use? | 0.5 | The code is licensed | | | | | | |
| | | | | 0.5 | License allows academic use | | | | | | |
| Interface | interface | availability | Does the tool have a graphical user interface? | 0.5 | The tool can be run using a graphical user interface, either locally or on a web server | 0 | 0 | 0 | 1 | 1 | 0 |
| | | | | 0.5 | The tool can be run through the command line or through a programming language | 1 | 1 | 0 | 1 | 1 | 1 |
| Function and object naming | naming | code_quality | Do the functions and objects have a consistent naming? | 0.67 | Functions/commands have well chosen names | 1 | 0 | 0 | 1 | 1 | 1 |
| | | | | 0.33 | Arguments/parameters have well chosen names | 1 | 1 | 0 | 1 | 1 | 1 |
| Code style | style | code_quality | Do the functions have a consistent style? | 0.5 | Code has a consistent style | 1 | 1 | 0 | 1 | 1 | 1 |
| | | | | 0.5 | Code follows (basic) good practices in the programming language of choice, for example PEP8 or the tidyverse style guide | 1 | 1 | 0 | 1 | 1 | 1 |
| Code duplication | duplication | code_quality | Is code frequently duplicated? | 1 | Duplicated code is minimal | 1 | 1 | 0 | 1 | 1 | 1 |
| Self-contained functions | pure_functions | code_quality | Does the code expose certain steps in the method as self-contained functions or commands? | 1 | The method is exposed to the user as self-contained functions or commands | 1 | 0 | 0 | 1 | 1 | 1 |
| Plotting | plotting | code_quality | Does the package allow plotting of (intermediate) results? | 1 | Plotting functions are provided for the final and/or intermediate results | 0 | 1 | 0 | 1 | 1 | 1 |
| Dummy proofing | dummy_proofing | code_quality | Does the package include dummy proofing? | 1 | Package contains dummy proofing, i.e. testing whether the parameters and data supplied by the user make sense and are useful | 0 | 0 | 0 | 0 | 0 | 0 |
| Unit testing | testing | code_assurance | Does the package include some testing? | 0.5 | Method is tested using unit tests | 0 | 0 | 0 | 0 | 0 | 0 |
| Unit testing | testing | code_assurance | Does the package include some testing? | 0.5 | Tests are run automatically using functionality from the programming language | 0 | 0 | 0 | 0 | 0 | 0 |
| Continuous integration | continuous_integration | code_assurance | Does the package include some continuous integration? | 1 | The method uses continuous integration, for example on Travis CI | 0 | 0 | 0 | 0 | 0 | 0 |
| Code | code_c | code_ass | Is the code coverage | 1 | The code coverage of the repository is assessed. | 0 | 0 | 0 | 0 | 0 | 0 |

| | | | | | | | | | | | |
|---|---|---|---|---|---|---|---|---|---|---|---|
| coverage | overage | urance | assessed? | 1 | What is the percentage of code coverage | 0 | 0 | 0 | 0 | 0 | 0 |
| Support | support | code_assurance | Does the package include a system to ask for support? | 0.5 | There is a support ticket system, for example on Github | 1 | 1 | 0 | 1 | 1 | 1 |
| | | | | 0.5 | The authors respond to tickets and issues are resolved within a reasonable time frame | 1 | 0 | 0 | 0 | 0 | 1 |
| Development model | development_model | code_assurance | Does the repository follow a development model, such as GitFlow? | 0.4 | The repository separates the development code from master code, for example using git master en developer branches | 1 | 0 | 0 | 0 | 0 | 0 |
| | | | | 0.4 | The repository has created releases, or several branches corresponding to major releases. | 1 | 0 | 0 | 0 | 0 | 1 |
| | | | | 0.2 | The repository has branches for the development of separate features. | 1 | 0 | 0 | 0 | 0 | 1 |
| Tutorial | tutorial | documentation | Is there a tutorial available for the method? Does this tutorial show everything the user needs? | 0.25 | A tutorial or vignette is available | 1 | 1 | 0 | 1 | 1 | 1 |
| | | | | 0.25 | The tutorial has example results | 1 | 0 | 0 | 1 | 1 | 0 |
| | | | | 0.25 | The tutorial has real example data | 1 | 1 | 0 | 1 | 1 | 0 |
| | | | | 0.25 | The tutorial showcases the method on several datasets (1=0, 2=0.5, >2=1) | 0 | 0 | 0 | 0 | 0 | 0 |
| Function documentation | documentation | documentation | Is the purpose and usage of each function documented? | 0.33 | The purpose and usage of functions/commands is documented | 1 | 0 | 0 | 0 | 0 | 0 |
| | | | | 0.33 | The parameters of functions/commands are documented | 1 | 0 | 0 | 1 | 1 | 0 |
| | | | | 0.33 | The output of functions/commands is documented | 0 | 0 | 0 | 0 | 0 | 0 |
| Inline documentation | inline_documentation | documentation | Is the code documented inline? | 1 | Inline documentation is present in the code | 1 | 0 | 0 | 0 | 0 | 0 |
| Parameter transparency | parameter_transparency | documentation | Are all important parameters available to the user? | 1 | All important parameters are exposed to the user | 1 | 0 | 0 | 1 | 1 | 1 |
| Unexpected output | unexpected_output | behaviour | Is unexpected output generated by the method? | 0.25 | No unexpected output messages are generated by the method | 1 | 0 | 0 | 0 | 0 | 0 |
| | | | | 0.25 | No unexpected files, folders or plots are generated | 1 | 1 | 0 | 1 | 1 | 0 |
| | | | | 0.5 | No unexpected warnings during runtime or compilation are generated | 0 | 0 | 0 | 0 | 0 | 0 |

| | | | | | | | | | | | |
|---|---|---|---|---|---|---|---|---|---|---|---|
| Relevant output | format | behaviour | Was postprocessing necessary to get the output of the method into a useful format? | 1 | The postprocessing necessary to extract the relevant output from the method is minimal (1), moderate (0.5) or extensive (0) | 1 | 0 | 0 | 1 | 1 | 1 |
| Publishing | publishing | paper | Is the method published? | 1 | The method is published | 0 | 1 | 1 | 1 | 1 | 1 |
| Peer review | peer_review | paper | Is the method published in a peer-reviewed journal? | 1 | The paper is published in a peer-reviewed journal | 0 | 1 | 0 | 1 | 0 | 1 |
| Evaluation on real data | evaluation | paper | Is the methods usefulness shown in the paper? | 0.5 | The paper shows the method's usefulness on several (1), one (0.25) or no real datasets. How many datasets was the model trained on | 1 | 0.25 | 1 | 1 | 0.25 | 0.25 |
| | | | | 0.5 | The paper quantifies the accuracy of the method | 0 | 1 | 1 | 1 | 1 | 1 |
| Evaluation of robustness | robustness | paper | Does the paper assess method robustness? | 1 | The paper assessed method robustness (to eg. noise, subsampling, parameter changes, stability) in one (0.5) or several (1) ways | 0 | 0 | 0 | 1 | 0 | 0 |
| Executing code | reproducibility | reproducibility | Did the code need significant adjustment to run | 2 | Able to run analysis pipeline with minimal/no changes to code | 0 | 0 | 0 | 1 | 1 | 0 |
| Domain expertise | domain_expertise | reproducibility | Does the method require domain expertise to run? | 1 | Experience in deep learning/Python/R is required (0), or not required (1) | 0 | 0 | 0 | 0 | 0 | 0 |
| Using models on new data | new_data | generalisability | Did the code require new code or code adjustments to predict new histology images | 1 | Method includes functionality to read and predict on new images | 0.5 | 0 | 0 | 0 | 0 | 0 |
